# A DRG genetic toolkit reveals molecular, morphological, and functional diversity of somatosensory neuron subtypes

**DOI:** 10.1101/2023.04.22.537932

**Authors:** Lijun Qi, Michael Iskols, David Shi, Pranav Reddy, Christopher Walker, Karina Lezgiyeva, Tiphaine Voisin, Mathias Pawlak, Vijay K. Kuchroo, Isaac Chiu, David D. Ginty, Nikhil Sharma

## Abstract

Mechanical and thermal stimuli acting on the skin are detected by morphologically and physiologically distinct sensory neurons of the dorsal root ganglia (DRG). Achieving a holistic view of how this diverse neuronal population relays sensory information from the skin to the central nervous system (CNS) has been challenging with existing tools. Here, we used transcriptomic datasets of the mouse DRG to guide development and curation of a genetic toolkit to interrogate transcriptionally defined DRG neuron subtypes. Morphological analysis revealed unique cutaneous axon arborization areas and branching patterns of each subtype. Physiological analysis showed that subtypes exhibit distinct thresholds and ranges of responses to mechanical and/or thermal stimuli. The somatosensory neuron toolbox thus enables comprehensive phenotyping of most principal sensory neuron subtypes. Moreover, our findings support a population coding scheme in which the activation thresholds of morphologically and physiologically distinct cutaneous DRG neuron subtypes tile multiple dimensions of stimulus space.

## Introduction

Sensory neurons of dorsal root ganglia (DRG) detect mechanical, thermal, and chemical stimuli acting on the body and transduce them into electrical impulses that are relayed to the central nervous system (CNS). DRG neurons exhibit a common morphological motif, with each having a peripheral axon that extends into the skin or an internal organ and a centrally projecting axon that terminates in the spinal cord. While this general architecture is shared, somatosensory neuron subtypes are distinguished by a wide range of molecular, morphological, and physiological properties underlying their selective responses to diverse environmental stimuli. A central question in somatosensory neurobiology is how primary sensory neuron subtypes contribute to the neural encoding of myriad stimuli acting on the body.

Early efforts to classify DRG neurons were based on their physiological response properties measured by nerve recordings. These foundational studies have inspired the current somatosensory neuron taxonomy. Focusing on cutaneous DRG neurons, those that are robustly activated by light, innocuous mechanical forces applied to the skin are termed low-threshold mechanoreceptors (LTMRs) and are subdivided into Aβ, Aδ, and C subtypes, which have fast, intermediate, and slow conduction velocities, respectively^1^. By contrast, DRG neurons with comparatively elevated mechanical force thresholds are termed high-threshold mechanoreceptors (HTMRs), and these neurons can exhibit either an Aδ or C conduction velocity^2^. Some subsets of mechanically sensitive Aδ or C fiber neurons also respond to thermal stimuli, and these neurons are often referred to as “polymodal nociceptors” ^3^ or, more specifically, A-and C-mechano-heat (MH), mechano-cold (MC), and mechano-heat-cold (MHC) neurons^2, 4, 5^. Alternatively, C fiber neurons may be sensitive to thermal stimuli but unresponsive to mechanical stimuli^6^, and these have been denoted C fiber heat (C-H) or cold (C-C) DRG neurons. Furthermore, small molecules, such as capsaicin, menthol, and histamine, can activate subsets of DRG neurons expressing relevant receptors^7–11^. In addition to the cutaneous innervation, a subset of DRG neurons send projections to internal organs instead of the skin, and many internal organs are thought to have some component of their neural innervation arising from DRGs^12, 13^. Finally, skeletal muscle and tendon innervating DRG neuron subtypes underlie proprioception^14^.

How do the physiological properties of DRG neuron subtypes overlay onto other features such as molecular profile, morphology, and central synaptic partners? Over the past two decades, the identification of marker genes and subsequent generation of mouse reporter alleles useful for labeling cutaneous DRG sensory neurons with defined properties has facilitated characterization of LTMRs and other subtypes^1, 15–34^. These molecular genetic tools have enabled a range of anatomical, physiological, functional, and behavioral analyses. Despite this, the properties and functions of a large number of DRG neuron subtypes identified using classic electrophysiological approaches remain difficult to explore due to a lack of labeling strategies. Thus, genetic labeling strategies for interrogating many of the principal DRG sensory neuron subtypes are needed to advance the field.

Our ability to explore the properties and functions of DRG sensory neurons has benefited by progress in genome-wide transcriptomics, in particular the large-scale sequencing of tens of thousands of single sensory neuron transcriptomes^28, 35, 36^. Collectively, these sequencing studies have revealed well over a dozen transcriptionally distinct subtypes of DRG sensory neurons and the genes they express. Here, we take advantage of previously reported^28^ and new large-scale transcriptomic datasets to generate a range of new mouse lines and assemble a near comprehensive genetic toolkit enabling comparisons across the principal transcriptionally defined subtypes of cutaneous DRG sensory neurons. We report that each transcriptionally defined subtype exhibits distinguishable morphological and physiological properties. Moreover, a comparative functional analysis of these populations revealed the contributions of each subtype to the encoding of the full physiological range of mechanical and thermal forces acting on the skin, supporting a role for population coding scheme in peripheral mechanosensation and thermosensation.

## Results

### A toolbox for genetic access to principal DRG neuron subtypes

To directly compare the morphological and physiological properties of the major classes of DRG neurons, we first performed a new large-scale single cell RNA sequencing (scRNA-seq) analysis of DRG neurons from 3-week-old mice with the goal of using this new information in conjunction with findings from a previously generated dataset^28^ to gain genetic access to each principal subtype. We obtained the transcriptomes from ∼40,000 DRG neurons, which were clustered and visualized by PCA/UMAP analysis (Figure 1A). Consistent with previous scRNA-seq taxonomic studies of DRG neurons^35–38^, the new dataset revealed at least 15 transcriptionally defined neuronal clusters corresponding to distinct somatosensory neuron subtypes based on previously identified marker genes. These are: Aβ RA-LTMRs, Aβ Field-LTMRs/Aβ SA1-LTMRs, Aδ-LTMRs, C-LTMRs, and CGRP^+^ neurons (containing at least seven transcriptionally discrete clusters), a PV^+^ cluster designated as proprioceptors, an MRGPRD^+^ cluster, a SST^+^ cluster and a TRPM8^+^ cluster (Figure 1A) ^15, 16, 19, 21, 25, 37, 39–50^.

**Figure 1.**
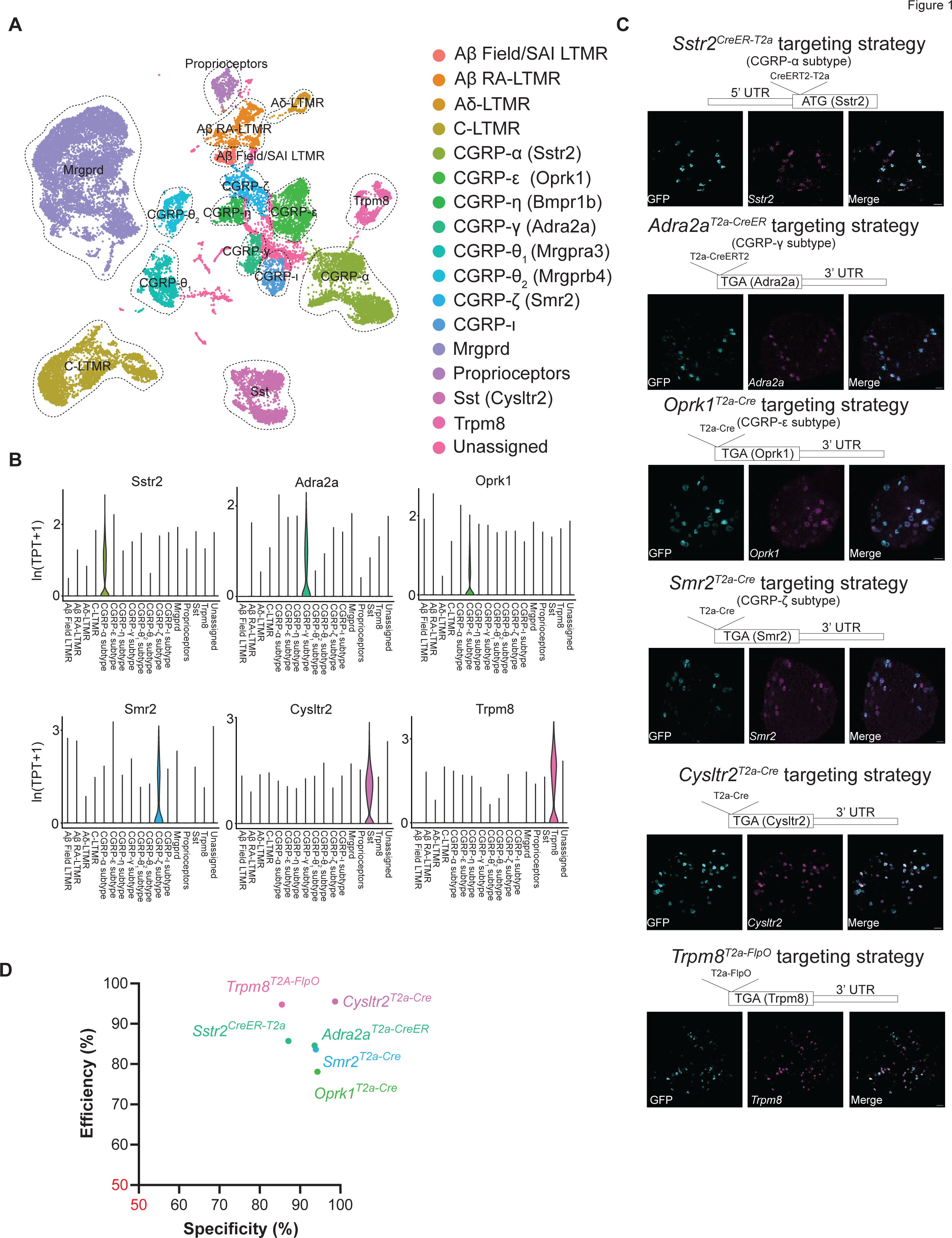
The transcriptional landscape of DRG sensory neurons informs the creation of new tools for genetic access to neuronal subtypes. **(A)** UMAP visualizations of DRG scRNA-seq data alongside putative sensory neuron subtype identities. **(B)** Violin plots displaying expression profiles of the marker genes used to generate the mouse recombinase lines. TPT: tags per ten thousand. **(C)** Targeting constructs and validation for the novel genetic tools using *in situ* hybridization, including *Sstr2^CreER-T^*^2a^ (to label CGRP-α), *Adra2a^T^*^2a–CreER^ (to label CGRP-γ), *Oprk1 ^T^*^2a–Cre^ (to label CGRP-ε), *Smr2^T^*^2a–Cre^ (to label CGRP-ζ), *Cysltr2^T^*^2a–Cre^ (to label CYSLTR2^+^/SST^+^ neurons) and *Trpm8^T^*^2a–FlpO^ (to label TRPM8^+^ neurons). *Smr2^T^*^2a–Cre^ mice were crossed to *R26^LSL-ReaChR^* mice; *Trpm8^T^*^2a–FlpO^ mice were crossed to *Avil^Cre^*; Ai195 (*TIGRE^LSL-jGCaMP^*^7s–FSF^) mice; the remainder were labeled using neonatal I.P. injections of an AAV carrying a Cre-dependent GFP reporter. Scale bar = 50 micron. **(D)** Summary of the specificity and efficiency of the novel genetic tools using the labeling strategies in (C). Specificity refers to the percentage of labeled cells that express the corresponding marker determined for all labeled DRG neurons; efficiency refers to the percentage of labeled cells that express the corresponding marker among all neurons that express the marker in the same section.

Using this DRG sensory neuron transcriptomic atlas, we sought to collate a mouse genetic toolkit that enables genetic access to each of the major transcriptionally defined DRG neuron subtypes, under the premise that each one is a morphologically, physiologically, and functionally distinct somatosensory neuron subtype. Clusters in the scRNA-seq atlas corresponding to the LTMR subtypes had previously validated genetic tools and labeling strategies. These are the C-LTMRs (labeled using *Th^2A-CreER^*) ^27^; Aδ-LTMRs (*TrkB^CreER^*) ^19^; Aβ RA-LTMRs (*Ret^CreER^*) ^17^; and Aβ SA1-LTMRs/Aβ Field-LTMRs (*TrkC^CreER^*; E12/postnatal Tamoxifen (TAM), respectively) ^15^ (Figure 1A, Table 1). In addition to the LTMRs, we noted that the cluster expressing MRGPRD is faithfully and selectively labeled using the *Mrgprd^CreER^*allele^30^, and the cluster representing proprioceptors is labeled using the *Pvalb^Cre^* allele^51^. However, strategies for selective genetic labeling of most of the remaining transcriptionally defined neuronal clusters in the scRNA-seq dataset, which represent nearly half of all clusters, are either not available or characterized to a lesser extent. Therefore, we generated new mouse lines for labeling transcriptionally distinct DRG neuron populations lacking existing tools, beginning with the clusters that express the peptide CGRP. We observed that CGRP^+^ subtypes were separable into at least seven transcriptionally distinct clusters, suggesting that there is considerable heterogeneity amongst neurons commonly and collectively referred to as “peptidergic nociceptors”. This finding of transcriptional heterogeneity of CGRP^+^ DRG neurons is consistent with prior immunohistochemical and electrophysiological analysis, indicating that CGRP^+^ neurons are comprised of functionally distinct subtypes^22–24, 37, 52, 53^. However, despite their implied nociceptive functions, few genetic tools useful for labeling and examining the properties and functions of transcriptionally distinct CGRP^+^ neuron subtypes exist. To name these uncharacterized CGRP^+^ subtypes, a Greek letter nomenclature had previously been used (Figure 1A, S1)^28^. We began by examining the two CGRP^+^ neuronal clusters that express the large caliber axon/myelination marker NF200 (*Nefh*) and the NGF receptor TrkA (*Ntrk1*) (Figure S1B), which are likely large diameter neurons with Aδ caliber axons. One of these clusters (CGRP-η) is robustly labeled using a recently generated allele, *Bmpr1b^T^*^2a–Cre^ ^28^, and we generated a new *Smr2^T^*^2a–Cre^ allele to label the second putative CGRP^+^ Aδ fiber population (CGRP-ζ) because differential gene expression analysis showed that the *Smr2* gene is preferentially expressed in this population (Figure 1B). Of the other CGRP^+^ clusters, which express low or undetectable levels of NFH (Figure S1B), and therefore are presumably C-fiber populations, we found that two preferentially expressed the *Mrgpra3* or *Mrgprb4* genes, which have been the basis for the *Mrgpra3^Cre^*and *Mrgprb4^Cre^* driver alleles (CGRP-θ_1_: Han *et al.* ^25^, CGRP-θ_2_: Vrontou *et al.* ^29^). The remaining CGRP^+^, NFH^-^, C-fiber neuron clusters did not have previously described genetic access strategies. Therefore, we used the differential gene expression analysis to identify genes enriched in these populations, and thus generated *Sstr2^CreER-T^*^2a^ (CGRP-α), *Oprk1^T^*^2a–Cre^ (CGRP-ε), and *Adra2a^T^*^2a–CreER^ (CGRP-γ) alleles to label three of them (Figure 1A-C). Lastly, we identified marker genes in the remaining two CGRP-negative populations and generated a *Cysltr2^T^*^2a–Cre^ allele and a *Trpm8^T^*^2a–FlpO^ allele (Figure 1A-1C) to label these clusters. We noted that the *Sst* gene is specifically expressed in the same cluster as *Cysltr2*^54, 55^; however, we observed inefficient recombination in the DRG using a previously published *Sst^IRES-Cre^*allele (Figure S1C) ^56^. Likewise, previously published *Trpm8* reporter alleles either used GFP^57, 58^ or were BAC transgenics^59^ with an uncertain degree of specificity. These considerations motivated the generation of the new *Cysltr2^T2A-Cre^* and *Trpm8^T^*^2a–FlpO^ knock-in alleles, for selective and versatile genetic labeling.

**Table 1.**
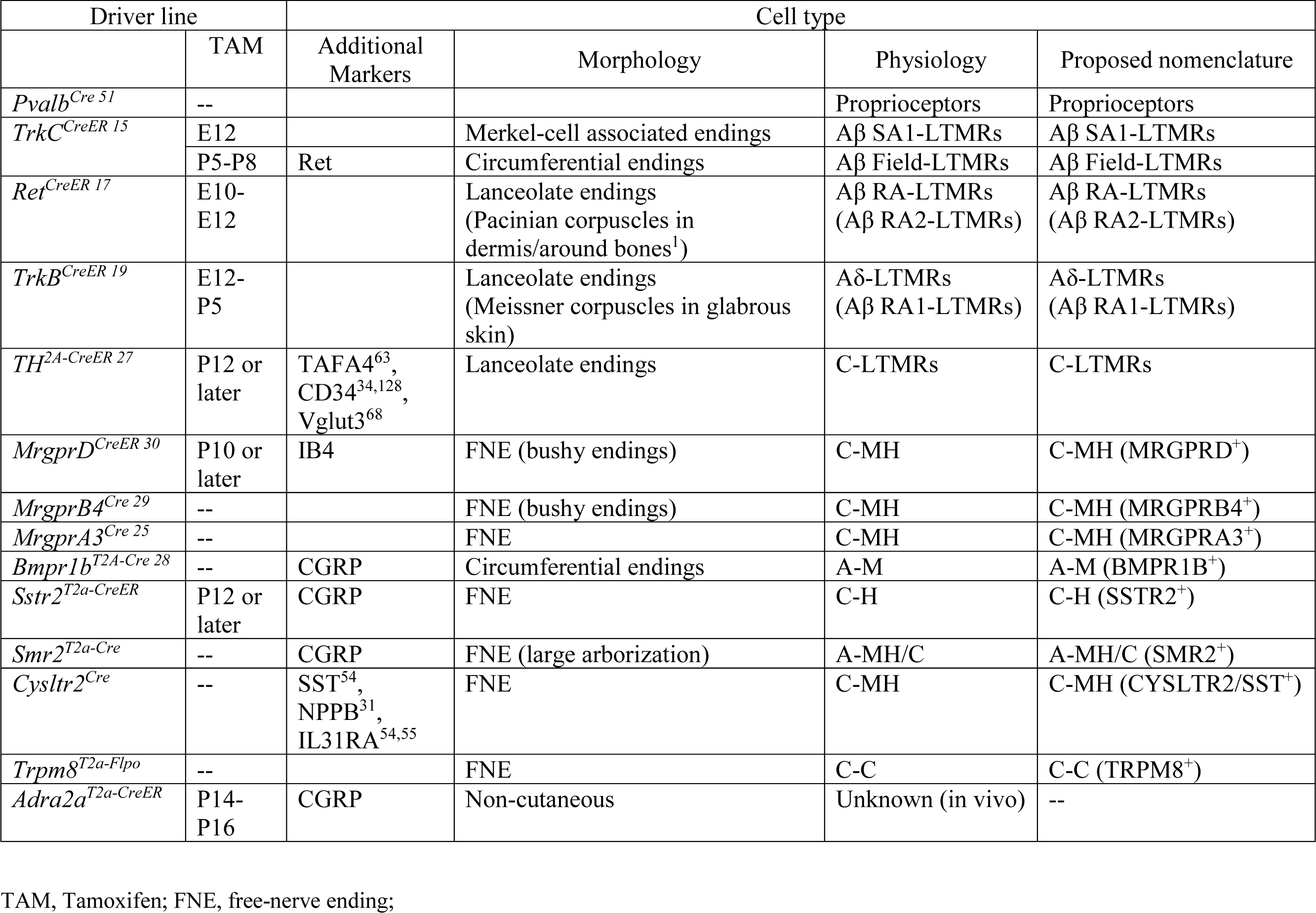
DRG somatosensory neuron genetic tools, physiological properties and nomenclature

Each of the new recombinase alleles (*Cysltr2^T^*^2a–Cre^, *Trpm8^T^*^2a–FlpO^, *Bmpr1b ^T2a-Cre^*, *Smr2^T2a-Cre^*, *Sstr2^CreER-T2a-^*, *Oprk1^T2a-Cre^*, and *Adra2a^T2a-CreER^*) was tested for specificity and efficiency of genetic labeling using double smRNA-FISH to detect expression of the gene used to generate the knock-in allele and the recombinase dependent reporter (Figure 1C, D). Consistent with specific, reliable genetic labeling, we found a high degree of overlap between the reporter gene labeled using the *Cysltr2^T2a-Cre^*, *Trpm8^T2a-FlpO^*, *Oprk1^T2a-Cre^*, *Smr2^T2a-Cre^*, *Sstr2^CreER-T2a^*, and *Adra2a^T2a-CreER^* mouse lines and the respective cell type specific marker genes. However, reporter gene expression in *Oprk1^T2a-Cre^* animals was not restricted to C-fiber CGRP^+^ DRG neurons (Figure 1A), as A-fiber (NFH^+^) or CGRP^-^ neurons were also labeled (Figure S1D, E), consistent with a previous study generating another Cre knock-in to the *Oprk1* locus using a different strategy^24, 60^, and therefore this allele was not used in subsequent analyses.

Interestingly, DRG neurons labeled using the *Adra2a^T2a-CreER^*(CGRP-γ) allele stood out in our initial analysis because they exhibited a striking difference in their prevalence across different axial levels (Figure S2A), which is consistent with our previous findings using *in situ* hybridization of transcripts selectively expressed in CGRP-γ neurons^28^. Because of this intriguing cellular distribution, we used *Adra2a^T2a-CreER^*mice and a Cre-dependent reporter allele (*Brn3a^cKOAP^* ^61^) expressed in the sensory neuron lineage to visualize the cell bodies and axonal processes of CGRP-γ neurons using whole mount alkaline phosphatase (AP) staining. This analysis revealed little to no axonal labeling in the skin of either the limbs or trunk in these mice (Figure S2B), but robust labeling of axonal terminals in internal organs, including the bladder (Figure S2C) and colon^62^. Thus, CGRP-γ neurons innervate non-cutaneous regions of the body. Notably, CGRP-γ neurons express the nicotinic acetylcholine receptor subunit alpha-3 (CHRNA3) gene^28^, showing similarity to *Chrna3^EGFP^* mice in which EGFP^+^ sensory fibers innervate internal organs and knees but not the skin^23^. However, compared to *Chrna3^EGFP^* mice, DRG neurons labeled in *Adra2a^T2a-CreER^* mice are more restricted to T7-L1 and L6-S1 ganglia, indicating the partial overlap of the labeling by these two lines (Figure S2A).

To further characterize the genetic strategies for labeling the transcriptionally distinct DRG neuron subtypes, we next sought to co-localize classical neurochemical markers and visualize the cell bodies of the genetically labeled neurons. This was achieved by expressing a cytosolic fluorescent reporter into each skin-innervating DRG neuron subtype using mice harboring one of the recombinase alleles and either a Cre-dependent reporter allele or neonatal intraperitoneal injection (IP) of an AAV carrying a Cre-dependent GFP construct (AAV-CAG-FLEX-GFP), with both approaches yielding similar results in DRG labeling. The reporter signals detected in DRG neuron cell bodies obtained from each of the recombinase mouse driver lines were colocalized with three neurochemical makers: CGRP, which labels subsets of small and large diameter DRG neurons; IB4, which labels several small diameter “nonpeptidergic” C-fiber neuron subtypes; and NFH, which labels medium and large diameter neurons that are lightly and heavily myelinated subtypes, respectively. This analysis revealed that the genetic approaches used to label clusters putatively assigned as Aβ and Aδ-fiber LTMRs have large diameter, NFH^+^ and CGRP^-^/IB4^-^ cell bodies, as expected. Also consistent with prior measurements, C-LTMRs are small diameter, CGRP^-^/IB4^-^ neurons (Figure S3A, C, D) ^16^. The genetically labeled MRGPRD^+^ neurons, CYSLTR2^+^ neurons, and TRPM8^+^ neurons exhibited small diameter cell bodies that are CGRP^-^/NFH^-^ (Figure S3A-D), also as expected^30, 54, 57^. Notably, while MRGPRD^+^ neurons were IB4^+^, consistent with previous reports^30^, CYSLTR2^+^ neurons and TRPM8^+^ neurons exhibited little overlap with IB4 (Figure S3A, C). Interestingly, while MRGPRA3^+^ and MRGPRB4^+^ populations were NFH^-^, as predicted^25, 63, 64^ (Figure S3B, C), both populations showed less overlap than expected with CGRP, which contrasts with the expression of CGRP transcripts observed by scRNA-seq (Figure S1B): we hypothesize that the mismatch in CGRP mRNA and protein in these subtypes is due to differences in translation. In the case of *Mrgprb4^Cre^*, which labels a large array of distinct DRG populations ^63^, expression is likely to be dynamic with multiple subtypes expressing Cre recombinase at different times throughout development, leading to genetic labeling of the sum of all populations that express *Mrgprb4* at any point in the life of the animal when crossed to a reporter line. As such, we use the designations MRGPRB4^SUM^ to refer to this population. The CGRP-α cluster neurons labeled using *Sstr2^CreER-T2a^* were CGRP^+^, but IB4^-^ and NFH^-^, and exhibited small-diameter cell bodies (Figure S3A-D). It is notable that despite the CGRP-α population being the most numerous CGRP^+^ subtype (Figure 1A), specific markers for this population have not been previously described. Lastly, the CGRP-η (BMPR1B^+^) and CGRP-ζ (SMR2^+^) populations were CGRP^+^, IB4^-^, and moderately NFH^+^, consistent with the likelihood that both populations are lightly myelinated with an Aδ conduction velocity (Figure S3A-D). These findings show that DRG populations labeled using the range of recombinase tools are relatively homogeneous with respect to their cell body diameters and neurochemical properties, which were predicted by the scRNA-Seq data.

Taken together, our expanded transcriptomic efforts combined with a series of newly generated mouse alleles (*Cysltr2^Cre^*, *Trpm8^T2a-FlpO^*, *Smr2^T2a-Cre^*, *Sstr2^CreER-T2a^*, *Adra2a^T2a-CreER^*, and *Bmpr1b^T2a-Cre^*) and previously reported alleles (*Th^T2A-CreER^*, *TrkB^CreER^*, *TrkC^CreER^*, *Ret^CreER^*, *Pvalb^Cre^*, *Mrgprd^CreER^*, *Mrgpra3^Cre^*, and *Mrgprb4^Cre^*) have led to the curation of a genetic resource useful for labeling, visualizing, and manipulating nearly all of the major transcriptionally distinct cutaneous DRG neuron populations (Table 1).

### Skin innervation patterns and morphological reconstructions of DRG sensory neuron subtypes

Genetic access to the range of transcriptionally distinct cutaneous DRG neuron subtypes allowed us to test the hypothesis that each subtype exhibits a distinguishable morphology. We next visualized the cutaneous endings of the labeled DRG neuron subtypes with single neuron resolution by performing sparse genetic labeling using the array of driver lines and whole-mount alkaline phosphatase (AP) staining of trunk hairy skin. Previous work has shown there is a morphological diversity of sensory nerve terminals in hairy skin^1^, including several hair-follicle associated endings (i.e. Merkel cell-associated endings, lanceolate endings, and circumferential endings) and free nerve endings. Our analysis showed that individual Aβ SAI-LTMRs innervated only one or two clusters of Merkel cells (touch domes), whereas individual Aβ RA-LTMRs, Aδ-LTMRs and C-LTMRs branched much more extensively, and to varying degrees, and formed lanceolate endings surrounding dozens of hair follicles (Figure 2A, E, F), consistent with previously reported observations^15, 65^. Also consistent with prior findings, individual Aβ field-LTMRs exhibited expansive morphological receptive fields and formed circumferential endings associated with many hair follicles (Figure 2B, E, F)^15^. Single-neuron analysis of MRGPRD^+^ neurons revealed a ‘bushy ending’ morphology, with densely packed clusters often embedding circumferential-like endings near hair follicles (Figure 2C, Figure S4A), in line with a previous report^30^. Most MRGPRB4^+^ neurons, selectively labeled by injection of an AAV expressing Cre-dependent AP at 3-4 weeks old, shared features of the “bushy ending” morphology but with larger arborizations compared to MRGPRD^+^ neurons^20, 66^, while MRGPRA3^+^ neurons exhibited branches that are less dense and an even larger terminal area (Figure 2C, E and G, Figure S4). Thus, three C-fiber populations, MRGPRD^+^, MRGPRB4^+^ and MRGPRA3^+^ neurons, exhibited related but quantitatively distinguishable morphologies (Figure S4D, E). In comparison, the “free-nerve endings” formed by individual CYSLTR2^+^ neurons and TRPM8^+^ neurons exhibited distinct branching patterns, with many fewer branches than the “bushy endings” formed by MRGPRD^+^ and MRGPRB4^+^ neurons (Figure 2C,G).

**Figure 2.**
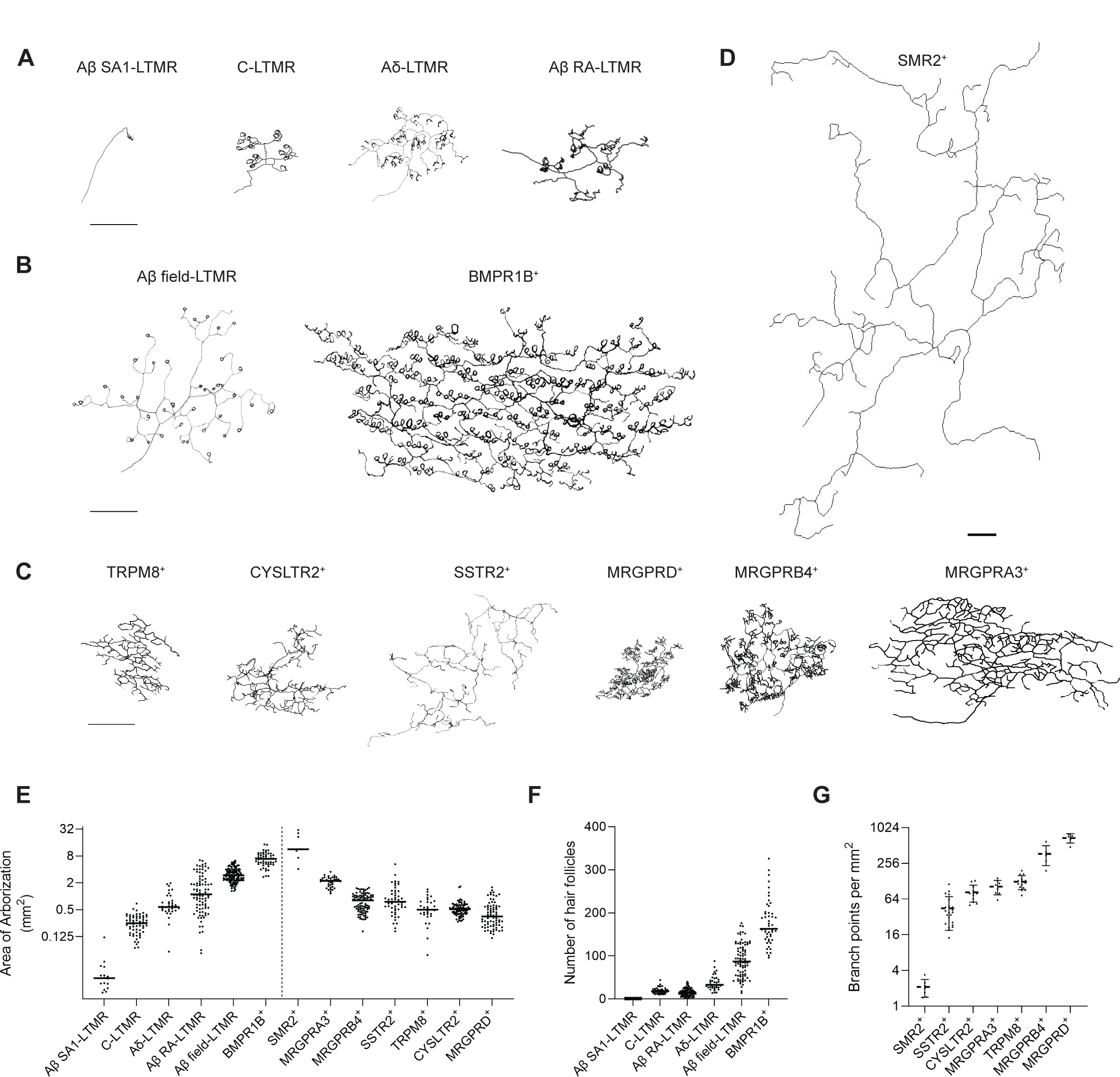
Morphological diversity of genetically labeled DRG subtypes revealed by sparse labeling. **(A)** Reconstructed examples of hairy skin whole mount AP staining of a single Aβ SAI-LTMR labeled using *TrkC^CreER^*; an Aβ RA-LTMR labeled using *Ret^CreER^*; an Aδ-LTMR labeled using *TrkB^CreER^* and a C-LTMR labeled using *TH*^2a–CreER^. All the driver lines in (A) are crossed to *Brn3a^cKOAP^*. **(B)** Reconstructed examples of an Aβ field-LTMR labeled using *TrkC*^CreER^, *Brn3a*^cKOAP^ and an CGRP-η neuron labeled using *Bmpr1b^Cre^* (AAV-CAG-FLEX-PLAP injection into hairy skin). **(C)** Reconstructed examples of free-nerve endings of individual TRPM8^+^, CYSLTR2^+^, SSTR2^+^, MRGPRD^+^, MRGPRB4^+^ and MRGPRA3^+^ neurons. See Methods for sparse labeling approaches. **(D)** Reconstructed examples of free nerve endings of a SMR2^+^ neuron. **(E)** Summary of the anatomical receptive field size of genetically labeled DRG subtypes. The dashed line separates the hair follicle-associated endings and “free-nerve” endings. The scale of the Y axis is log2. **(F)** Summary of the number of hair follicles innervated by individual neurons of the different subtypes. **(G)** Summary of branching density of free-nerve ending neurons. All scale bars 500 µm (A-D). The data for Aβ SAI-LTMRs, Aβ RA-LTMRs, and Aβ field-LTMRs are replotted from Bai *et al.*^15^. The example Aβ SAI-LTMR and Aβ field-LTMR neurons were reconstructed from data reported in Bai *et al.* ^15^.

We next assessed the single-neuron morphologies of the transcriptionally distinct clusters of CGRP^+^ neuronal subtypes. Individual SSTR2^+^ neurons exhibited relatively few branches and widely spaced axon terminals across the arborization area (Figure 2C, G, Figure S4A). Individual BMPR1B^+^ neurons, in contrast, formed circumferential endings associated with hair follicles at a quantity double that of Aβ field-LTMRs, and this corresponded to larger anatomical receptive fields than Aβ field-LTMRs (Figure 2B, E, F). Thus, the circumferential endings of BMPR1B^+^ neurons and Aβ field-LTMRs have large innervation areas, with BMPR1B^+^ neurons being the largest among the hair follicle-associated afferents (Figure 2E, F). Finally, SMR2^+^ neurons exhibited expansive morphological receptive fields with very sparse branches (Figure 2D, E, G, Figure S4A-C). These remarkably expansive SMR2^+^ neurons formed the largest anatomical receptive field among the entire cohort of DRG neuron subtypes, reaching sizes up to 30mm^2^ (Figure 2E). The distinct morphologies of BMPR1B^+^ and SMR2^+^ neurons revealed an underappreciated morphological diversity of Aδ CGRP^+^ DRG neurons.

To complement the whole-mount AP visualization of individual neurons across the major cutaneous DRG neuron subtypes, we expressed a fluorescent reporter into each subtype and performed immunostaining analyses using antibodies against tdTomato or GFP with co-staining of CGRP in hairy skin samples obtained from 3-4-week-old mice. This analysis allowed us to localize the cutaneous axon terminals in the skin and assess the extent to which they terminate in association with hair follicles or in the dermis, or whether they penetrate the epidermis. Consistent with previous observations and the single neuron reconstructions, Aβ RA-LTMRs, Aδ-LTMRs, and C-LTMRs all form lanceolate endings associated with the hair follicles (Figure S5B) ^1, 16, 19^. On the other hand, MRGPRD^+^, CYSLTR2^+^, MRGPRB4^SUM^, MRPGRA3^+^ and TRPM8^+^ neurons terminated as free nerve endings within the epidermis (Figure S5). For the CGRP^+^ populations, SSTR2^+^ neurons formed free nerve endings in the epidermis, whereas the two Aδ caliber CGRP^+^ populations showed remarkably distinct morphologies: BMPR1B^+^ neuron terminals formed circumferential endings that wrap around the hair follicles (Figure S5), consistent with previous reports ^22, 28^, whereas SMR2^+^ neurons formed a dense plexus of free nerve endings that penetrated the epidermis (Figure S5). The finding that SMR2^+^ neurons are a major population of A-fiber CGRP^+^ fibers that penetrate the epidermis is notable and stands in contrast to the commonly held assumption that CGRP^+^ neurons that innervate the epidermis are predominantly slow conducting C-fiber neurons.

Thus, across the transcriptionally distinct subtypes there exists a remarkably large range of terminal morphologies and branching patterns; each genetically defined cutaneous DRG neuron subtype exhibits a unique, distinguishable morphology. Moreover, of the transcriptionally distinct cutaneous DRG neuron subtypes that innervate hairy skin, six form axonal endings that associate with hair follicles and seven form free nerve endings that penetrate the epidermis.

### A comparison of the central projection patterns across DRG sensory neuron subtypes

To compare and contrast the central axonal projection patterns across the transcriptionally defined neuronal subtypes, spinal cords of genetically labeled mice were harvested and triple immunolabelling was performed using antibodies against the reporter to visualize the genetically labeled axons, CGRP to label CGRP^+^ axons, and IB4 to label non-peptidergic C fiber axons that terminate within lamina IIi. Beginning superficially, TRPM8^+^ neurons were observed to terminate within the most dorsal region of the dorsal horn, immediately superficial to CGRP^+^ fibers of lamina I (Figure 3A, C). Terminals of SSTR2^+^ neurons, BMPR1B^+^ neurons, and SMR2^+^ neurons were primarily located in lamina I and IIo, with some SMR2^+^ terminals also observed in deeper lamina (Figure 3A, C). Continuing deeper in the dorsal horn, the CYSLTR2^+^, MRGPRA3^+^, MRGPRB4^SUM^ and MRGPRD^+^ populations terminated primarily in lamina IIi (Figure 3A-C). Then, deeper in the dorsal horn, C-LTMR terminals were concentrated in lamina IIiv, Aδ-LTMR axons were observed in lamina IIiv and lamina III, and Aβ RA-LTMRs terminated in lamina III-IV (Figure 3B, C), consistent with prior reports ^27^. Therefore, the central terminals of transcriptionally distinct DRG neuron subtypes elaborate within defined domains of the spinal cord.

**Figure 3.**
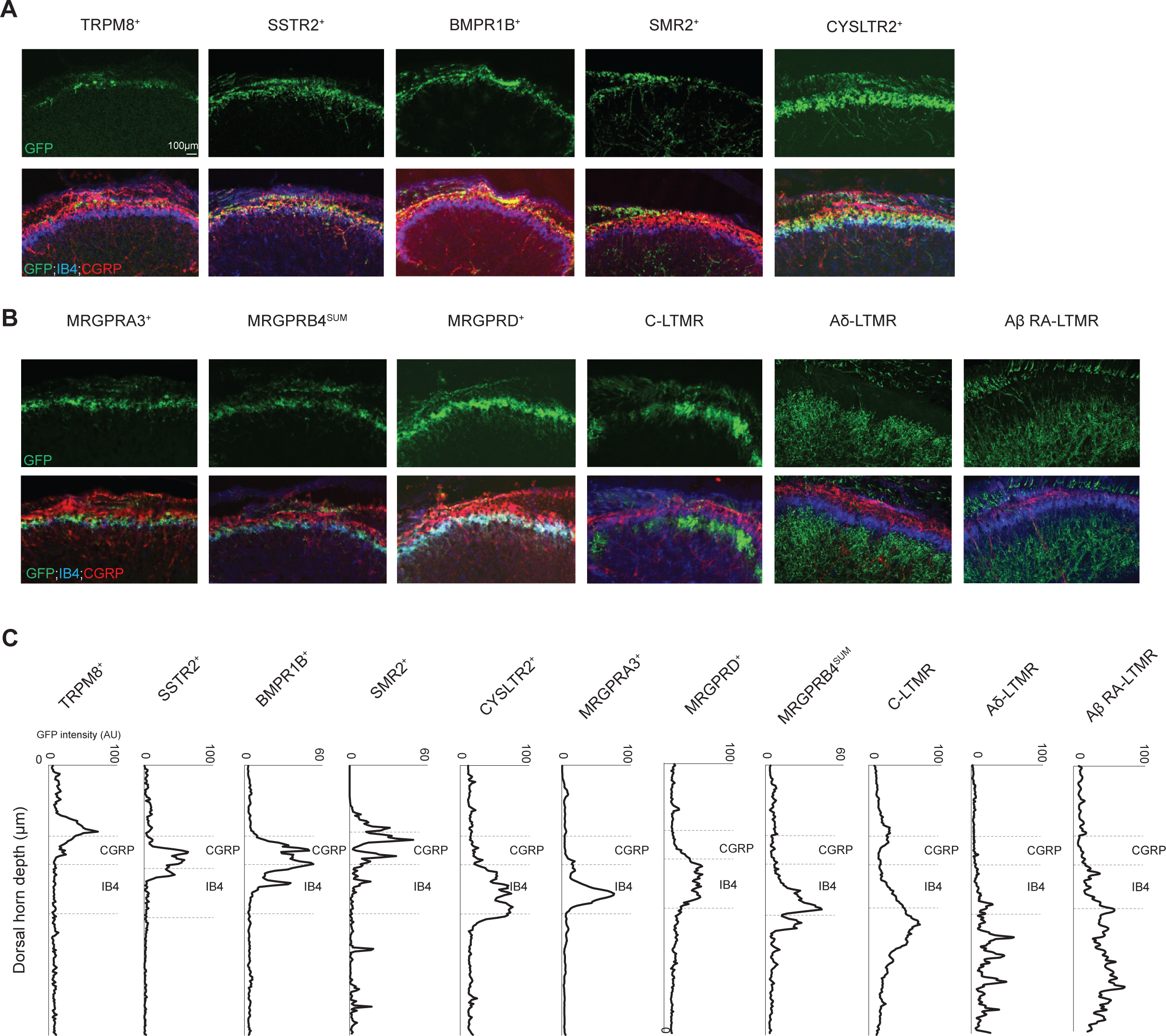
Characterization of the spinal cord terminals of genetically labeled cutaneous DRG subtypes. **(A and B)** Representative immunostaining images of the central terminals in the lumbar spinal cord (∼L3-L5) from mice labelled using new sensory neuron subtype genetic tools (A) and existing driver lines (B). The sections were co-stained with CGRP and IB4. **(C)** Quantification of the depth of the axon terminals in spinal cord dorsal horn.

Taken together, DRG neurons genetically labeled using the curated genetic toolkit exhibited uniform cutaneous morphologies, branching patterns, cell body sizes, expression of classical markers, and central termination zones. These findings underscore the selectivity and utility of the cutaneous DRG neuron mouse genetic toolkit (Table 1).

### Indentation force space is tiled by the mechanical thresholds of at least 10 DRG neuron subtypes

We next used the DRG genetic toolkit to determine the degree to which different sensory stimuli activate the transcriptionally defined sensory neuron populations. To achieve this, *in vivo* calcium imaging of L4 DRG neurons was performed using the Cre or FlpO mouse lines in conjunction with GCaMP reporters (Figure 4A). We started by examining mechanical sensitivity across the cutaneous DRG subtypes, using step indentations of the skin. For this, a 200 µm diameter indenter tip was used to deliver forces ranging from 1 to 75mN to the receptive fields of neurons, identified by mechanical or electrical probing to the thigh hairy skin (Figure 4A). These measurements allowed for a quantitative comparison of indentation response thresholds and adaption properties across the subtypes.

**Figure 4.**
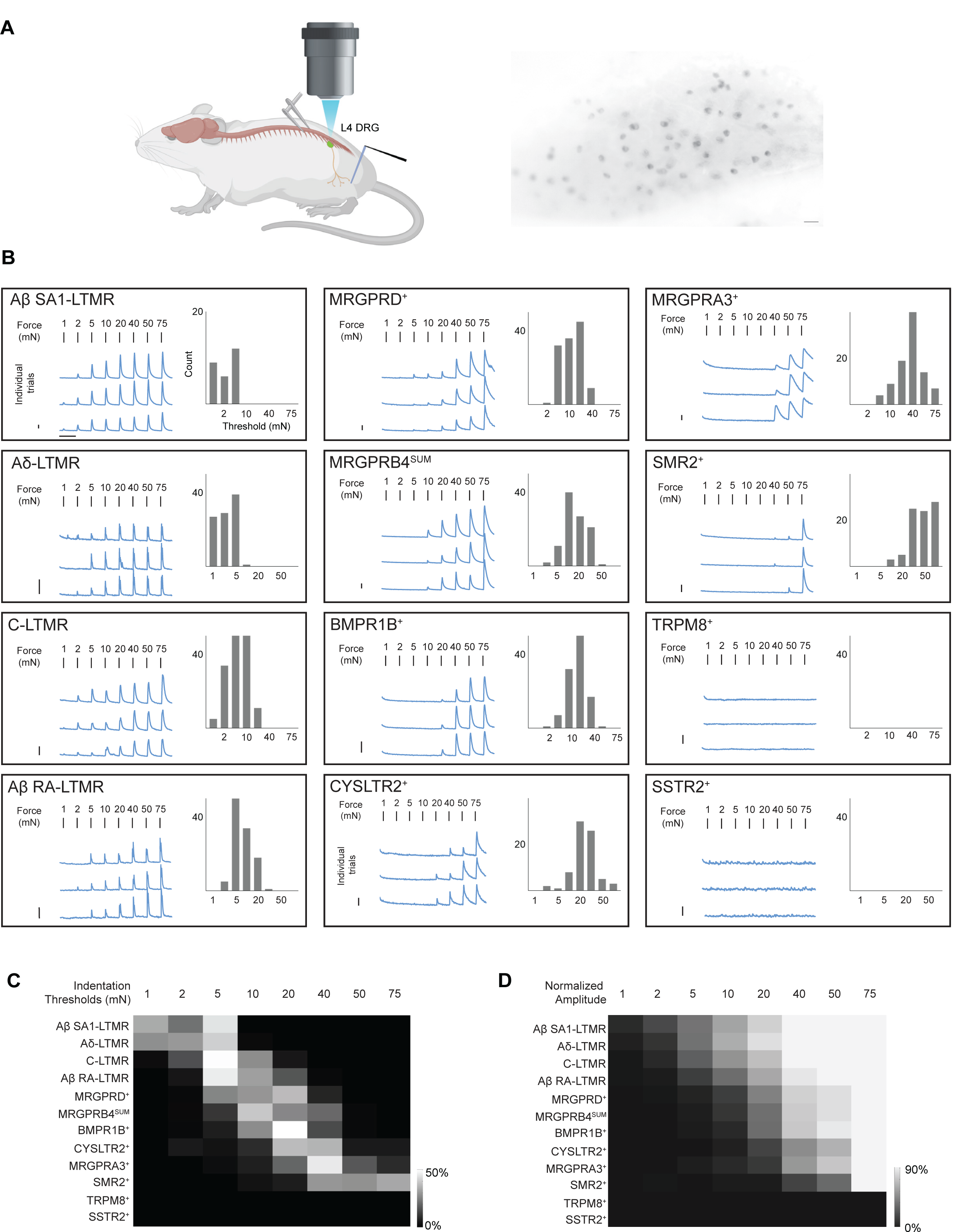
Indentation force space is tiled by the different mechanical thresholds of DRG neuron subtypes. **((A)** Left: Schematic of *in vivo* DRG calcium imaging and application of indentation. Right: Representative field of view from *Th^2A-CreER^*; Ai148 (intensity of baseline fluorescence, scale bar 50µm). To express GCaMP in distinct DRG subtypes, *Bmpr1b^T^*^2a–Cre^ was crossed to Ai95; *MrgprA3^Cre^*was crossed to Ai96; *Trpm8^T^*^2a–FlpO^ was crossed to *Avil^Cre^;* Ai195; the other recombinase lines were crossed to Ai148. **(B)** Representative calcium signals and threshold distribution of each DRG neuron subtype responding to 0.5-second step indentations. Left in each box: Traces from a total of three trials for the same example neuron; The scale bars are 20% ΔF/F for Y and 10s for X. Right in each box: Number of traces with certain threshold. **(C)** Diverse response profiles to cooling. The cooling-activated C-LTMRs exhibit transient response to temperature decrease. TRPM8^+^ neurons show 3 types of responses: cooling-activated (top), cooling-activated and warmth-inhibited (middle), and cooling-and-heat activated (bottom) responses. **(D)** Diverse response profiles to warmth or heat. MRGPRB4^SUM^, CYSLTR2^+^ and MRGPRA3^+^ neurons have two types of responses: neurons responding to relative increase (top) and only to absolute temperature (bottom). The SSTR2^+^ and MRGPRD^+^ neurons respond to absolute warmth or heat. **(E)** SMR2^+^ neurons exhibit response to heat and/or cold. Scale bars in C-E are 20% ΔF/F for Y and 10s for X. Individual traces are shown in gray, and the average trace is shown in blue. Dashed vertical lines indicate the change of temperature.

We found that Aβ RA-LTMRs, Aβ SAI-LTMRs, Aδ-LTMRs, and C-LTMRs innervating hairy skin all exhibited a low-threshold activation pattern, as expected. Specifically, Aβ SAI-LTMRs, Aδ-LTMRs and C-LTMRs showed exquisite mechanical sensitivity, with most neurons responding at ∼1-5 mN indentation force (Figure 4B, C). Aβ RA-LTMRs had slightly higher thresholds, in general, but most responded in the low-threshold range (≤10mN) (Figure 4B, C). By comparing the amplitude of peak calcium signals for each force step (normalized to the maximum force delivered), each of these four LTMR subtypes exhibited graded increases in responses in the low force range and plateaued between 20mN and 40mN (Figure 4D, Figure S6C), consistent with the notion that most LTMRs preferentially encode skin indentation forces in the low intensity range.

By contrast, MRGPRD^+^ and MRGPRB4^SUM^ neurons showed graded responses over a broader range of mechanical forces, with most neurons exhibiting force thresholds between 5 to 20mN, which is in the low to medium force intensity range (Figure 4B-D). The BMPR1B^+^ neurons exhibited slightly higher force thresholds, most around 20mN. The thresholds of CYSLTR2^+^ neurons and MRGPRA3^+^ neurons were higher still, in the 20-40mN range. SMR2^+^ neurons exhibited the highest force thresholds, responding primarily to forces greater than 40mN (Figure 4B, C). Lastly, TRPM8^+^ neurons and SSTR2^+^ neurons were unresponsive to mechanical indentations at all forces tested (Figure 4B-D), despite their responsivity to other stimuli (Figure 5A). Therefore, considering their mechanosensitivity and/or preferential tuning to intense mechanical stimuli (Figure 4C, D, Figure S6C), at least six DRG neuron populations, the MRGPRD^+^, MRGPRB4^SUM^, BMPR1B^+^, CYSLTR2^+^, MRGPRA3^+^ and SMR2^+^ neurons, fall into the “HTMR” category, or the “nociceptor” category based on the classical nociceptor definition^2, 3^, while TRPM8^+^ and SSTR2^+^ neurons are insensitive to mechanical indentation of the skin.

**Figure 5.**
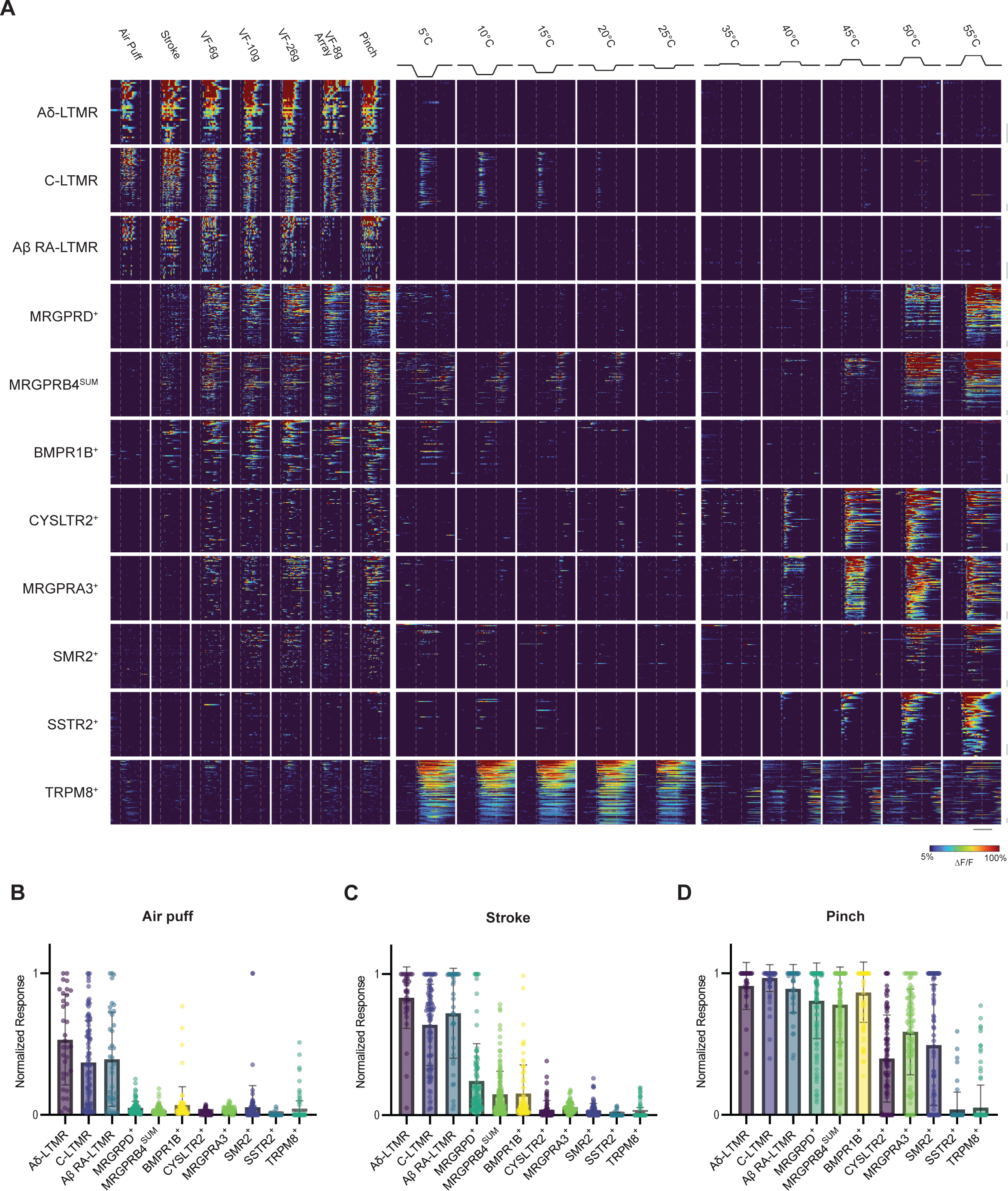
Responses to diverse mechanical and thermal stimuli by distinct DRG neuron subtypes. **(A)** Heatmaps of calcium signals for individual neurons stimulated with a range of mechanical and thermal stimuli. VF-6g, VF-10g and VF-26g refer to 6-gram, 10-gram and 26-gram Von Frey filaments. VF-8 gram Array refers to an array (5*5) of custom made 8-gram Von Frey hair mounted on a holder of the same size as Peltier device (See Methods). The baseline of temperature was 32^°^C. Each row of the heatmap represents responses of an individual neuron. The vertical scale bars in the right refer to 10 neurons. The horizontal scale bar is 20 seconds. **(B-D)** Summaries of the normalized response amplitudes for air puff (B), stroke (C) and pinch (D) stimuli. The amplitudes (ΔF/F) were normalized to the maximum responses to all stimuli of the same neurons.

We also asked whether the subtypes exhibit transient or sustained responses to static indentation of the skin. To accomplish this, a complementary series of mechanical indentation experiments were done in which the duration of skin indentation time was extended to three seconds (Figure S6A). We observed that nearly all Aδ-LTMRs responded exclusively during both the onset and offset of indentations, but not during the sustained indentation period, and most Aβ RA-LTMRs (29/36) also responded during the onset and offset of step indentations with a subset (7/36) responding only at stimulus onset (Figure S6B). In contrast, Aβ SAI-LTMRs exhibited sustained calcium signals with large amplitudes for the duration of the indentation step, as expected. We further observed that C-LTMRs had an intermediate rate of adaption, with large responses observed at both the onset and offset of the stimulus and a slow decrease of the calcium signal during the sustained phase (Figure S6B), also in line with previous measurements ^16^. By contrast, each of the six HTMR subtypes (MRGPRD^+^, MRGPRB4^SUM^, BMPR1B^+^, CYSLTR2^+^, MRGPRA3^+^ and SMR2^+^ neurons) exhibited sustained calcium responses during the entire three second step indentation period (Figure S6B), suggesting that each of these subtypes is slowly adapting to sustained mechanical stimuli.

We next determined whether the genetically labelled DRG neuron subtypes respond to an array of additional cutaneous stimuli (Figure 5). For this, the skin was stimulated using an air puff (1 PSI), gentle stroke using a cotton tip, pinch with forceps, and indentation with Von Frey filaments. Thermal stimuli were also applied using a custom-made Peltier device contacting the same area of skin that received the mechanical stimuli.

While both air puff and stroking across the skin deflect hairs, the stroke stimulus also gently indents the skin with forces <50mN. Therefore, air puff and stroke both represent innocuous stimuli, albeit with distinct characteristics. In contrast, pinch of the skin, using a round rubber pad (0.25cm^2^) mounted on forceps and delivered to a large area of skin with a force in the range of 3-5 N, is a noxious, high force mechanical stimulus. We tested three LTMR subtypes (Aβ RA-LTMRs, Aδ-LTMRs, and C-LTMRs) in this analysis and observed that both air puff and cotton swab stroke elicited robust responses (Figure 5A-C), consistent with the step indentation studies (Figure 4, S6), showing that these populations are highly sensitive to light mechanical stimuli, as expected. Consistent with the idea that LTMRs saturate in the innocuous range (Figure 4D, Figure S6C) ^1^, these same populations also generally reached maximum activation by stroke (Figure 5C), with pinch showing no greater level of activation.

As with the LTMRs, three HTMR populations, BMPR1B^+^ neurons, MRGPRB4^SUM^ neurons, and MRGPRD^+^ neurons, were also activated by skin stroke. However, in contrast to the LTMRs, these three populations exhibited a higher degree of activation by pinch (Figure 5C, D). Thus, while BMPR1B^+^ neurons, MRGPRB4^SUM^ neurons, and MRGPRD^+^ neurons can be activated by innocuous forces, they reach peak responsivity at noxious forces, reflecting their wide dynamic range of mechanosensitivity (Figure 4C, D). The CYSLTR2^+^ neurons, MRGPRA3^+^ neurons, and SMR2^+^ neurons showed much higher activation by pinch than stroke, but their responses to pinch were smaller than their responses to thermal stimuli, on average (Figure 5A, C, D, Figure 7). Lastly, most TRPM8^+^ and SSTR2^+^ neurons exhibited little or no response to any of the mechanical stimuli, including pinch and Von Frey filaments of up to 26 grams (Figure 5A, C), which is consistent with the step indentation analysis (Figure 4B-D, Figure S6B) and indicates that these two populations are mechano-insensitive.

In summary, ten of the transcriptionally distinct cutaneous DRG neuron subtypes tested here encode stimulus intensity across a range of innocuous to noxious mechanical forces, with each subtype exhibiting a unique force threshold. Two other DRG neuron subtypes showed weak or no responsivity to any of the mechanical stimuli tested. Also, when considered as a collective, the transcriptionally distinct DRG mechanoreceptor subtypes tile the physiological range of indentation force thresholds (Figure 4C). These observations are consistent with a population coding scheme, in which an increasing number of LTMR and HTMR subtypes are recruited as skin indentation intensity increases, with individual subtypes specialized at encoding within a distinct force range.

### Temperature is represented as absolute or relative by different sensory neuron subtypes

Following delivery of mechanical stimuli, the same region of skin was contacted with a custom-built Peltier system, and quantitative pulses of thermal stimuli were delivered after the neurons had adapted to the light mechanical contact of the Peltier, more than two minutes later. A series of rapid temperature steps were delivered, starting from physiological skin surface temperature (32°C) and then changing (5°C s^-1^) and being held (20 seconds) at a target temperature ranging from 5°C to 55°C. The temperature was reverted to physiological skin surface temperature, again at a rate of 5°C s^-1^ (Figure S6D). The temperature steps were delivered sequentially, beginning with innocuous and then into the noxious range, with alternating presentations of heat and cold stimuli.

Cooling of the skin activated both TRPM8^+^ neurons and C-LTMRs; however, the responses of these two populations were highly distinct. While TRPM8^+^ neurons exhibited heightened activity for the duration of the cooling epoch (20s) and exquisite sensitivity to cool temperatures (Figure 5A, 6A, 6C), the C-LTMRs responded transiently and exclusively during the ramping periods and only when temperature was decreasing (Figure 5A, 6C). Moreover, the C-LTMR responses to cooling were considerably smaller in amplitude compared to their responses to mechanical stimuli (Figure 5A, 7A). These observations are consistent with prior observations that C-LTMRs are activated during cooling of the skin ^16, 67, 68^. We also observed that the TRPM8^+^ and C-LTMR populations responded to temperature decreases from warm/heat to baseline (32^°^C), although with diminished amplitudes compared to cooling below 32^°^C (Figure 6C). Thus, both TRPM8^+^ and C-LTMR populations are sensitive to decreases in skin surface temperatures; TRPM8^+^ neurons are sensitive to absolute cooling whereas C-LTMRs respond when temperature is actively decreasing. Intriguingly, around 40% of TRPM8^+^ neurons exhibited inhibition by warmth (Figure 6C), suggesting that this population is tonically active at physiological skin temperatures, consistent with recent reports ^6, 69^. We also note that a single, discrete DRG neuron subtype dedicated to encoding noxious cold temperatures (<15^°^C) was not identified. Rather, we observed that responses to noxious cold temperatures are distributed across subsets of multiple neuronal subtypes (Figure 5A), including 19% of SMR2^+^ neurons (Figure 6E), 8% of TRPM8^+^ neurons, 10% of BMPR1B^+^ neurons, 11% of MRGPRD^+^ neurons, 8% of MRGPRB4^SUM^ neurons and 9% of SSTR2^+^ neurons.

**Figure 6.**
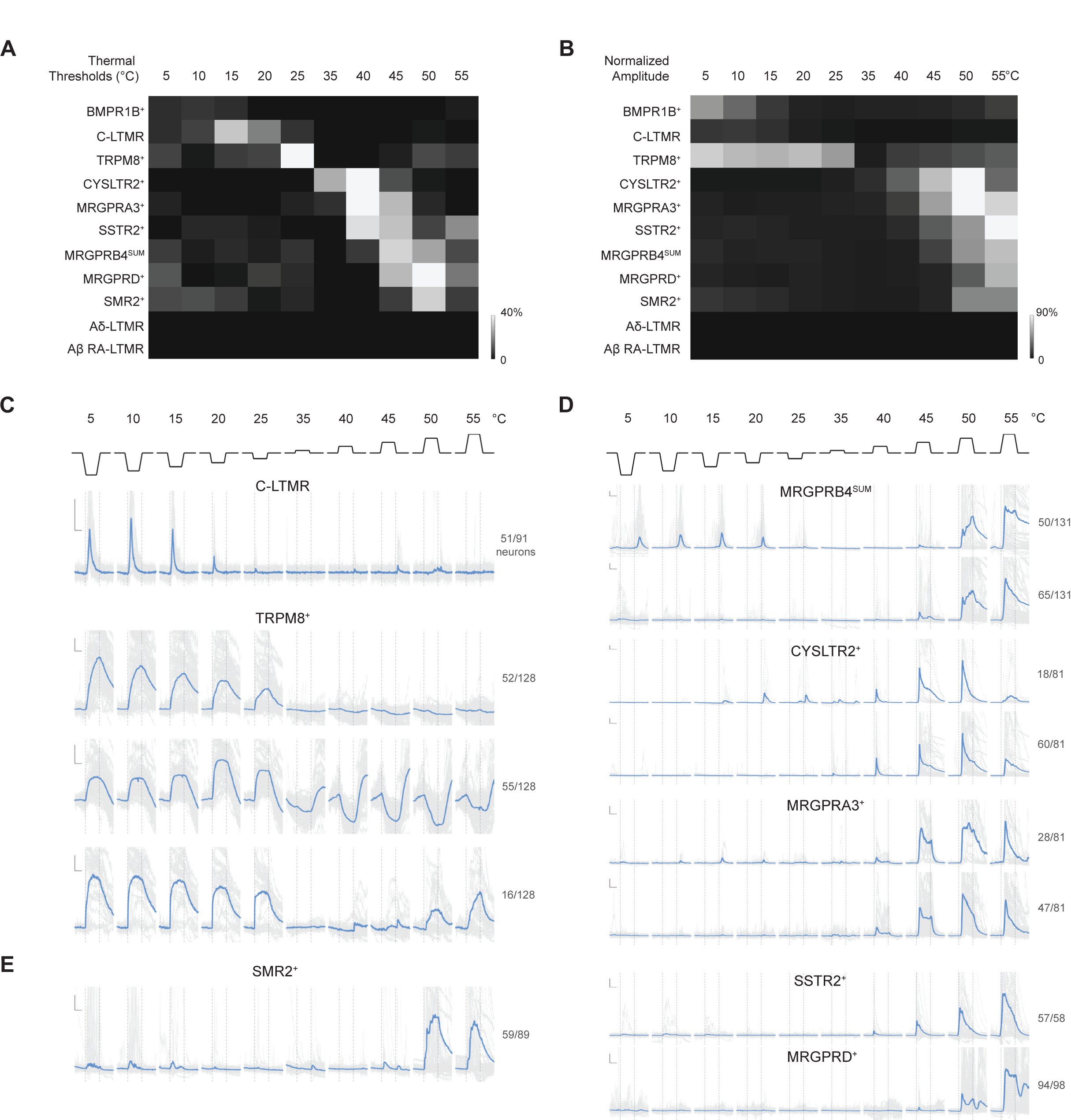
Temperature is reported as absolute or relative by distinct sensory neuron subtypes. **(A)** Summary of thermal thresholds for the DRG subtypes. The percentage of neurons with a particular thermal threshold (among all neurons of each subtype) is represented by the brightness of the heatmap. Due to the lack of thermal responses in Aβ RA-LTMRs and Aδ-LTMRs, the percentages at all temperatures are set to zero. **(B)** Summary of normalized response amplitudes for the DRG subtypes. The amplitude (ΔF/F) of thermal sensitive neurons from each subtype is normalized to the maximum response from all stimuli of the same neurons, then averaged within each subtype. The amplitudes of Aβ RA-LTMRs and Aδ-LTMRs were set to zero due to the same reason in (A). **(C)** C-LTMRs exhibit transient response to decreases of temperature. Note that responses to changes from warm to baseline (32^°^C) are considerably smaller than responses from baseline to cold temperature. **(D)** Subsets of CYSLTR2^+^, MRGPRB4^SUM^, and MRGPRA3^+^ neurons encode relative increases in temperature, while responses to changes from cold to baseline is smaller than from baseline to warm temperatures. **(E)** A subset of TRPM8^+^ neurons were inhibited by warmth, while showing rebound during changes from warm/heat to baseline. **(F)** A different subset of TRPM8^+^ neurons showed robust responses to both heating and cooling, and some also responded to temperature decreases, from 40^°^C to baseline.

**Figure 7.**
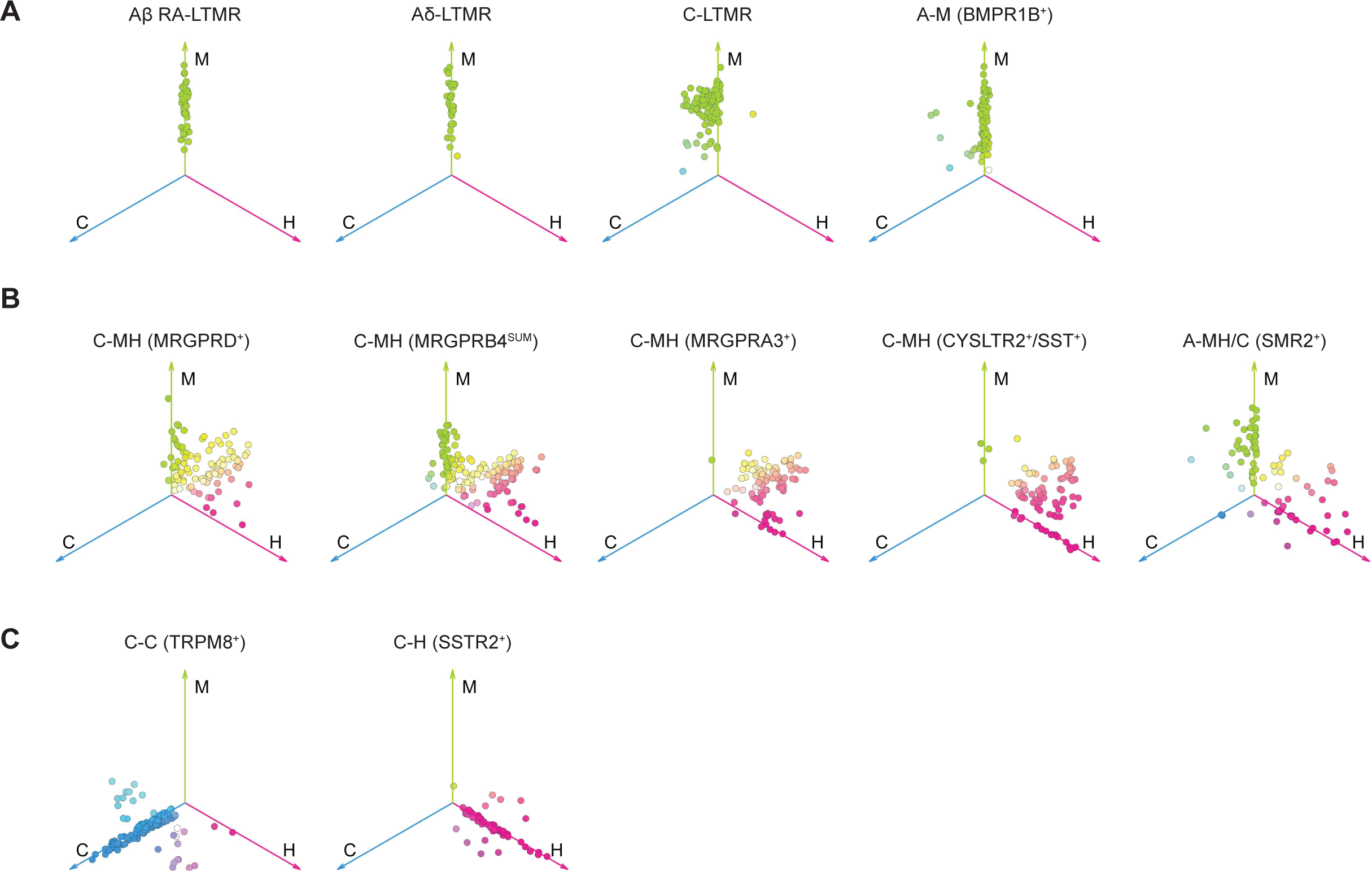
Polymodality of DRG neuron subtypes. **(A-C)** Distributions of tuning preferences of each DRG subtype in the polymodality space. Each dot represents a neuron. The location of each dot was determined by the summation of the magnitude of the vector for responses to mechanical (M), heat (H) or cold (C) stimuli. The color of each dot represents the neuron’s relative preference to the three modalities, thus a white dot near the center indicated that the neuron is tuned to all three modalities. The subtypes with strong preference to mechanical stimuli are plotted in **(A)**; subtypes with polymodal responses are plotted in **(B)**; subtypes with predominantly thermal responses are plotted in **(C)**.

Seven of the transcriptionally defined DRG neuron populations responded to elevations in skin temperature: SSTR2^+^, MRGPRD^+^, MRGPRB4^SUM^, MRGPRA3^+^, CYSLTR2^+^, SMR2^+^ neurons and a subgroup of TRPM8^+^ population (Figure 6A-E). The MRGPRD^+^, MRGPRB4^SUM^, and SMR2^+^ neurons were most reliably activated by noxious heat (≥45^°^C), while many SSTR2^+^ neurons and MRGPRA3^+^ neurons had thresholds of ∼40^°^C (Figure 5A, 6A); neurons in these populations exhibited sustained activation throughout the duration of the thermal stimulation period (Figure 5A). The neurons most sensitive to innocuous warm temperatures (35-40^°^C) were the CYSLTR2^+^ neurons (Figure 6A). Interestingly, subsets of CYSLTR2^+^ (33%), MRGPRB4^SUM^ (38%) and MRGPRA3^+^ (35%) neurons were activated transiently during increases in temperature (Figure 6D), especially from cold to baseline, suggesting that these neurons are activated during the ramping phase of temperature elevation in addition to an absolute temperature.

These findings are consistent with a multipronged strategy for the encoding of thermal stimuli; individual DRG neuron subtypes encode either a decrease or increase in skin temperature. Moreover, within these two broad categories, subtypes are responsive to either transitions in temperature or absolute temperatures of the skin surface, with neurons in the latter category exhibiting distinct thresholds and sustained responses.

### Polymodality of transcriptionally distinct DRG neuron subtypes

By comparing responses to mechanical and thermal stimuli in the same neuronal populations, we were also able to quantitatively assess the extent of polymodality of individual neurons within each genetically labeled population by comparing the maximal responses to different modalities^70^. This analysis revealed that virtually all Aβ RA-LTMRs and Aδ-LTMRs and the majority of BMPR1B^+^ neurons are tuned to mechanical but not thermal stimuli, while C-LTMRs are sensitive to cooling but most strongly activated by mechanical stimuli (Figure 7A). The majority of MRGPRD^+^, MRGPRB4^SUM^, MRGPRA3^+^, CYSLTR2^+^, and SMR2^+^ neurons are polymodal, sensitive to both thermal and mechanical stimuli (Figure 7B) ^25, 71–73^. In contrast, the majority of SSTR2^+^ and TRPM8^+^ neurons are sensitive to thermal, but not mechanical stimuli (Figure 7C).

Finally, these functional analyses enable alignment of the classical physiology-based descriptions of neuronal populations^2^ with the transcriptionally distinct neuronal subtypes. Therefore, the classical physiology-based nomenclature used for the LTMRs (C-LTMRs; Aδ-LTMRs; Aβ RAI-LTMRs; Aβ RAII-LTMRs; Aβ SA-LTMRs; Aβ field-LTMRs) can be extended to the other subtypes of cutaneous DRG neurons. The classic nomenclature is based on mechanical and thermal responsivity measurements, where the first term is designated C or A, referring to C-and A-fiber conduction velocity, and the second term designation(s) are M, C, and H, which refer to mechanical, cold, and heat responsivity, respectively. Thus, for the C-fiber types: MRGPRD^+^ neurons are C-MH (MRGPRD^+^); CYSLTR2^+^ neurons are C-MH (CYSLTR2^+^/SST^+^); MRGPRB4^+^ neurons are C-MH (MRGPRB4^SUM^); TRPM8^+^ neurons are C-C (TRPM8^+^); SSTR2^+^ neurons are C-H (SSTR2^+^); and MRGPRA3^+^ neurons are C-MH (MRGPRA3^+^). For the A-fiber HTMR subtypes: BMPR1B^+^ neurons are A-M (BMPR1B^+^) and SMR2^+^ neurons are A-MH/C (SMR2^+^) (Table 1).

## Discussion

Defining the properties and functions of primary DRG sensory neurons is fundamental for understanding somatosensation. Accomplishing this requires phenotypic analysis of the principal DRG sensory neuron subtypes, across multiple levels of analysis ^74^. Here, we used large scale transcriptomic measurements of the mouse DRG to guide the assembly of an array of mouse lines for transcriptionally distinct DRG neuron subtypes. We used this resource of mouse lines to compare the morphological and physiological properties across the majority of transcriptionally defined cutaneous DRG neuron populations. Our findings reveal a remarkable degree of morphological and physiological diversity of at least 15 principal, transcriptionally distinct DRG neuron subtypes. The near comprehensive DRG sensory neuron toolkit will enable deep phenotypic and functional analyses of the neuronal building blocks of somatosensation.

### Genetic access to the principal DRG neuron subtypes

Genetic access to cell types has propelled the field of neuroscience forward by allowing interrogation of defined components of neural circuitry. To understand how somatosensory information is detected and relayed from the skin to the CNS, it is essential to define the morphological and physiological properties, synaptic connections, and functions of the primary sensory neuron subtypes. This has been difficult to achieve because identifying definitive markers of several if not most DRG neuron subtypes has been a challenge. Indeed, several commonly used marker genes, such as *Trpv1*, *Calca*, *Scn10a* (Na_V_1.8), and *Piezo2*, are expressed in multiple transcriptionally, morphologically, and physiologically distinct neuronal subtypes, and their expression patterns may change during development. Therefore, analyses using driver lines based on these and other genes inadvertently lead to multiple, potentially functionally orthogonal, populations being analyzed or manipulated. By contrast, identifying stable markers of terminally differentiated sensory neuron subtypes requires efforts that must include examination of anatomical and physiological properties to ensure that relatively homogenous populations of neurons are being labeled for analysis. For primary somatosensory neurons, we contend that transcriptomics coupled with morphological and physiological analyses enable resolution of the principal somatosensory neuron types. Nevertheless, it should be acknowledged that defining properties of subtypes may extend beyond the resolution of transcriptomics and genetics because some defining features of sensory neurons may also be shaped by the skin regions and cutaneous end organs within which their axons terminate.

Transcriptomic analyses, including new measurements reported herein, have revealed more than 15 transcriptionally distinct DRG neuron populations, and we have exploited these transcriptomic data to generate and curate a collection of genetic tools to investigate the major DRG neuron populations. Genetic access to several DRG neuron classes or subtypes, including C-LTMRs, Aδ-LTMRs, Aβ RA-LTMRs, Aβ field-LTMRs, Aβ SA-LTMRs, and the broad proprioceptor population, had been previously achieved^15–17, 19, 27, 30, 51^, and further confirmed using the genomics and functional analyses in the present study. The CGRP^+^ DRG neurons, long known to be physiologically and morphologically heterogeneous ^22, 37, 52^, were lacking genetic tools and, consequently, discerning the properties and functions of C-fiber and Aδ-fiber CGRP^+^ neuron subtypes had been a challenge. Resolving the morphological and physiological diversity of CGRP^+^ neuron subtypes is of particular interest because, as a broad population, CGRP^+^ neurons play important roles in nociception^22, 75^. The present study reports new tools for accessing CGRP^+^ neuronal subtypes and new insights into the properties and functions of the diverse CGRP^+^ populations. For example, CGRP^+^ epidermal-penetrating free nerve endings are comprised of intermingled Aδ-fibers (A-MH/C (SMR2^+^) neurons) and C-fibers (C-H (SSTR2^+^) neurons). Moreover, the CGRP^+^ A-MH/C (SMR2^+^) neurons with free nerve endings appear to be the only subtype with a response threshold in the noxious range, suggesting that these neurons are uniquely poised to rapidly alert the CNS to the presence of damaging mechanical or thermal stimuli acting on the skin. It is also noteworthy that at least one CGRP^+^ subtype, the CGRP-γ neurons, labeled using the *Adra2a^T2a-CreER^* allele, uniquely targets internal organs of the body and not the skin. Therefore, the assembled genetic tools that enable interrogation of each of the transcriptionally distinct CGRP^+^ subtypes, as well as other populations, will enable further assessment of how noxious stimuli acting on the skin and internal organs are encoded by primary sensory neurons and relayed to the CNS to initiate pain perception.

Several notable questions remain. First, at least one major DRG neuron subtype is not represented in our genetic toolkit: the morphological and physiological properties of the CGRP-ε population remain unknown, in part because our attempts to gain selective genetic access to this population have so far failed, and there may be additional CGRP^+^ subtypes (i.e., CGRP-ι, Figure 1A) yet to be resolved. Second, tools for Aβ field-LTMRs^15^, Aβ SA1-LTMRs^15^, and Aβ RA2-LTMRs (neurons that innervate Pacinian corpuscles^17^) exist but they are limited in terms of selectivity and ease of use, and thus these subtypes were not included in some or all experiments of the present study. Third, some DRG neuron populations are inevitably missing from our analysis; this may include subtypes that are relatively rare, such as Aβ RA2-LTMRs, or enriched in a subset of axial levels, as was observed for CGRP-γ (Adra2a^+^) neurons, or otherwise lowly abundant in the transcriptomic dataset ^26, 34^. Consistent with this, a previous study using sparse random labeling of DRG neurons revealed a neuron type exhibiting “large arbors with free endings (LA-FE)” ^76^, and this morphology was not observed in our morphological analysis using the genetic tools described herein. The LA-FEs exhibit a similar arborization area to that observed for SMR2^+^ neurons (Figure 2D, E), but with a highly distinct branching pattern (Figure 2G). It is also plausible and perhaps likely that some cutaneous DRG neuron subtypes can be further subdivided beyond what is presented here. Indeed, proprioceptors, labelled with *Pvalb^Cre^* based strategies can be subdivided into at least five transcriptionally, morphologically, and functionally distinct subtypes^77–79^. Related to this, we observed that some populations exhibited heterogeneous responses to mechanical and thermal stimuli (Figure 5A, Figure 7), including responses to noxious cold, which may represent further cell type specializations or subdivisions not demarcated in this study through transcriptomics and genetic labeling. As such, further subdivisions may arise from exceedingly difficult to detect transcriptional differences, rendering gene expression differences between cell type variants too small to exploit with existing mouse genetic approaches. Furthermore, the characterizations of the DRG neuron subtypes in this study are limited to healthy animals, and it would be fascinating to explore how the molecular, morphological, and physiological properties of these neuron subtypes change during pathological states^23, 59, 63, 71, 80–87^. Finally, we note that our analyses focused on hairy skin, which covers most of the mouse body and may host the most diverse cohort of DRG neuron types. Characterization of the genetically defined neuron subtypes defined here with respect to other skin regions, including glabrous skin of the paws and the external genitalia, as well as internal organs, will likely be fruitful and may lead to the identification of additional physiological and morphologically distinct DRG sensory neuron subtypes.

### Molecular basis of DRG neuron response properties

To relate the molecular basis of sensory reactivity to the observed physiological properties across the distinct DRG neuron subtypes, we compared the mRNA abundance of well-known mechanically or thermally activated ion channels across the neuronal subtypes using the single cell transcriptomic dataset (Figure S7). It is notable that all mechanosensitive DRG neuron subtypes exhibit moderate to high levels of Piezo2 expression^34, 82, 88–90^, while the mechano-insensitive subtypes, C-H (SSTR2^+^) and C-C (TRPM8^+^), exhibit minimal or undetectable expression of Piezo2. However, in addition to Piezo2, other factors are likely to contribute to the distinct mechanical thresholds, adaptation properties, and responsivity to hair deflection and vibratory stimuli of the distinct LTMR and HTMR subtypes. Additional determinants of mechano-responsivity include the axonal structures and other cellular constituents of mechanosensory end organs^1^, skin arborization patterns (Figure 2), the ion channel determinants of intrinsic excitability^37^, modulation by non-neuronal cells^65, 91–96^, expression of Piezo1^97^, and other potential mechanosensitive molecules and auxiliary proteins^98–101^.

On the other hand, thermosensitivity may be explained, at least in part, by unique combinations of TRP channels expressed in the transcriptionally distinct DRG neuron subtypes. The warm-sensitive neuronal subtypes, with thermal thresholds in the 35-40^°^C range, including C-MH (CYSLTR2^+^/SST^+^), C-MH (MRGPRA3^+^) and C-H (SSTR2^+^) neurons, express high levels of TRPV1 (Figure S7B), which contributes to warmth encoding ^59, 102–104^. C-MH (CYSLTR2^+^/SST^+^) neurons also express TRPM2 (Figure S7), another warm-sensing molecule^105^. Additionally, TRPM3, TRPV2 and TRPA1 are expressed in both C-MH (MRGPRD^+^) and C-MH (MRGPRB4^SUM^) neurons (Figure S7), potentially contributing to their heat sensitivity ^106–108^. TRPM8^+^ neurons uniquely express high levels of TRPM8^49, 50, 58, 109^, and a subset also express TRPV1 (Figure S7B), which may explain the heat sensitivity of some TRPM8^+^ neurons (Figure 6C). Curiously, C-LTMRs do not express detectable levels of known thermal TRP channels, and thus how these neurons respond to cooling the skin remains unclear. Furthermore, the diverse thermal response dynamics and the encoding of relative temperature (Figure 6C-D) may involve mechanisms beyond employing TRP channels expressed in DRG neurons^110–112^.

### Distinct DRG neuron types tile the spectrum of mechanical and thermal stimuli

By systematically comparing indentation force thresholds of genetically labeled DRG neurons, we found that thresholds form a graded distribution across the subtypes, with no well-defined boundary between LTMR and HTMR subtypes (Figure 4C). However, a clearer distinction between LTMR and HTMR subtypes could be elucidated when examining response saturation. Indeed, while activation of LTMR subtypes mostly saturated within the innocuous force range, as expected, at least five HTMR subtypes displayed modest activation by innocuous forces and comparatively enhanced activation by noxious forces, such as pinch (Figure 4D, 5C, 5D). These observations prompt us to propose a need for revising the classical, dichotomous view of LTMRs and “mechano-nociceptors” being two distinct groups of cell types responsible for non-overlapping functions.

Although it may be tempting to subsume HTMR subtypes under the broader “nociceptor” category, we suggest that this nomenclature is not ideal. The classical use of the term nociceptor refers to a neuron that encodes stimulus intensity into the noxious range, regardless of whether it exhibits a low threshold for spiking ^3^. However, this classical definition does not account for emerging evidence showing that activation of HTMRs does not necessarily lead to the perception of pain or a nocifensive response ^2, 71, 113^. Indeed, microneurography recordings in conscious human subjects revealed that mechanically evoked C-MH fibers can discharge even up to 10 Hz without an accompanying painful percept ^114^. Moreover, recent work in mice has indicated that optogenetic activation of neurons labeled by the *Mrgprd^Cre^* allele, which includes neurons that we classify here as C-MH (MRGPRD^+^) neurons, did not lead to place aversion in a place preference test ^71^, and optogenetic activation of C-MH (MRGPRD^+^) neurons did not trigger a “pain-like” behavior evaluated using a mouse pain scale ^113^. Similarly, C-MH (MRGPRB4^SUM^) neurons are implicated in conveying positive valence associated with touch ^29, 63^. These observations, combined with our current analysis indicating that innocuous forces activate most HTMR subtypes, albeit to a lesser extent than noxious forces, support the view that most HTMR subtypes contribute to the encoding of innocuous forces acting on the skin, rather than exclusively contributing to the perception of pain. We do, however, note that the A-MH/C (SMR2^+^) population of HTMRs represents an exception to this heuristic, as this population is activated by intense, or noxious mechanical and thermal stimuli, while exhibiting little to no observable activation in the innocuous stimulus range (Figure 4C, D, Figure 5A-D). Therefore, A-MH/C (SMR2^+^) neurons may be the only principal DRG neuron subtype exclusively tuned to noxious mechanical stimuli.

Related to this, certain DRG neuron subtypes are implicated in dimensions of somatosensation not tested here, such as chemical signaling, and some of these neuronal subtypes have been designated as pruriceptors (itch receptors). Indeed, selective activation of C-MH (CYSLTR2^+^/SST^+^) or C-MH (MRGPRA3^+^) neurons can evoke scratching behaviors^10, 11, 25, 31, 64, 97, 115^, and these subtypes express distinct receptors (Histamine H1 receptor, CYSLTR2, MRGPRA3, IL31RA etc.) ^54, 55, 116, 117^ rendering them sensitive to compounds capable of eliciting the perception of itch. It follows that chemical stimuli lead to combinatorial patterns of somatosensory neuron subtype activation not observed when compared to mechanical or thermal stimuli. It is therefore likely that different percepts arise when distinct combinations of subtypes are co-activated. In further support of this idea, it is noteworthy that both ‘pruriceptor’ populations are activated by indenting or warming of the skin, which do not necessarily evoke scratching behaviors. We therefore favor the view that a behavioral response or a particular percept arises from the combinatorial nature of the cohort of sensory neuron subtypes activated by a given stimulus, the convergence and integration of these signals in the spinal cord and brainstem, and the neural computations made by higher-order CNS neurons and circuits^7, 118, 119^. These deductions lead us to favor a taxonomy of primary somatosensory neurons that is based on physiological properties rather than any specific percept or behavior such as those implied by the terms “nociceptor” or “pruriceptor”.

By examining the physiological profiles across sensory modalities, we found that at least five of the genetically labeled DRG neuron populations exhibit polymodal responses: these are the C-MH (MRGPRD^+^) neurons ^71–73^; C-MH (CYSLTR2^+^/SST^+^) neurons; C-MH (MRGPRA3^+^) neurons ^25^; C-MH (MRGPRB4^SUM^) neurons, and A-MH (SMR2^+^) neurons. While C-LTMRs also respond to rapid cooling ^16, 67, 68^, we continue to refer to these neurons as “C-LTMRs” because their responses to light touch are more prominent than to cooling and they lack responses to absolute cool or cold temperature. Moreover, two types of C fiber neurons encode thermal but not mechanical stimulation of hairy skin: the majority of C-C (TRPM8^+^) neurons and C-H (SSTR2^+^) neurons respond to cold and warm temperatures, respectively, but not mechanical stimuli. Thus, in total, at least 12 morphologically and physiologically distinct DRG neuron subtypes encode mechanical forces (at least six LTMRs, the responses of two of these—Aβ field-LTMRs and Aβ RA2-LTMRs—were not studied here, and at least six HTMRs) and no fewer than eight morphologically and physiologically distinct subtypes encode thermal stimuli acting on the skin, with six exhibiting polymodality. Interestingly, mechanical force space is tiled by indentation force thresholds of the LTMR and HTMR subtypes. In parallel with this, each of the thermally responsive DRG neuron subtypes encodes thermal stimuli within a unique temperature range, and thus thermal space is also tiled by the response thresholds of thermally responsive subtypes. Thus, distinct cohorts of DRG neuron subtypes are recruited as the force of indentation increases or as the temperature of a thermal stimulus increases or decreases. This general inference may hold true across other “dimensions” of somatosensation. Indeed, for vibration tuning, low (1-10 Hz), medium (10-200 Hz), and high (50-1000 Hz) frequency vibrations of the skin are encoded by different mechanoreceptor subtypes^1^. Therefore, the somatosensory system is comprised of a large number (>15) of physiologically distinct primary sensory neuron subtypes with distinct activation thresholds for temperature and across different dimensions of mechanical stimulus space and chemical space, and thus combinatorial population and rate codes signify where within a particular dimensional space a stimulus resides.

## EXPERIMENTAL MODELS AND SUBJECT DETAILS

All experiments performed in this study were approved by the Institutional Animal Care and Use Committee (IACUC) of Harvard Medical School and Columbia University. Male and female mice of mixed genetic backgrounds were used for these studies.

### Mouse lines

Five of the new mouse lines (*Smr2^T2a-Cre^*, *Sstr2^CreER-T2a^*, *Adra2a^T2a-CreER^*, *Trpm8^T2a-FlpO^*, *Oprk1^T2a-Cre^*) were generated at the Gene Targeting and Transgenics Facility at Janelia Research Campus using standard homologous recombination techniques in hybrid mouse embryonic stem (ES) cells. These mouse lines were generated by knocking the T2a-(recombinase) cassette directly upstream of the stop codon in the last annotated exon of each indicated gene. *Sstr2* targeting was the exception as there were multiple annotated splice variants, therefore in this case, a (recombinase)-T2a cassette was knocked in directly downstream of the coding exon containing the methionine start codon for Sstr2. This design strategy was chosen to minimize potential deleterious consequences associated with deleting the gene. Chimeras were generated by blastocyst injection and germline transmission was confirmed by standard tail genotyping PCR. The neo selection cassette was left intact and each line was overtly phenotypically normal, even when bred as homozygous carriers of the knock-in allele, consistent with the introduction of the T2a-recombinase cassette minimally perturbing endogenous gene expression.

To generate *Cysltr2^T2a-Cre^*, a guide RNA (GAATTTCAAAGCTCGATTAA) was designed specific for the 3’ region of the murine Cysltr2 locus close to the STOP codon using the Zhang lab guide design tool (crispr.mit.edu, replaced by zlab.bio/guide-design-resources). sgRNA was generated by *in vitro* transcription using the MEGAshortscript T7 Transcription Kit from Invitrogen (cat. # AM1354) from a template carrying a T7 promoter, guide RNA and an RNA scaffold (based on Mali et al. ^120^) encoding the sgRNA. The Cas9 mRNA was generated using the mMESSAGE mMACHINE T7 Transcription Kit from Invitrogen (cat. # AM1344) from a template carrying a T7 promoter and a sequence encoding Cas9. Both, the sgRNA and Cas9 mRNA were purified using the MEGAclear Transcription Clean-Up Kit (cat. # AM1908). An embryo injection mix was prepared with the following components: (1) DNA donor molecule carrying a Cysltr2-2A-Cre recombinase targeting construct (100ng/ul) (2) sgRNA (112ng/ul) (3) Cas9 mRNA (100ng/ul). The DNA donor molecule was sequenced beforehand to confirm sequence fidelity. Embryos on the C57Bl/6 genetic background were injected. A chimeric founder mouse was identified by PCR interrogating the left and right integration sites of the targeting construct with the following primer combinations: MP319 and MP320 (left junction) yielding an 846bp product and MP302-MP317 (right junction) yielding a 554bp product. PCR products were TOPO cloned and sequenced to confirm faithful integration of the targeting construct. To investigate the possibility of unwanted, off-target activity of the injected CRISPR/Cas9 mix, we performed off-target analysis on the top 10 predicted off-targets and one additional off-target located within a gene (off-target 20). Specifically, we designed PCR primers around these predicted sites and sequenced the PCR products. Our analysis indicated that no edit occurred in this set of highest probability off-targets.

### Single-cell RNA-seq

The dissection strategy used was essentially as previously described ^28^. Briefly, animals were sacrificed at approximately postnatal day 21 and spinal columns rapidly removed and placed on ice. Individual DRG with central and peripheral nerves attached were removed from all axial levels and placed into ice-cold DMEM:F12 (1:1) supplemented with 1% pen/strep and 12.5 mM D-glucose. A fine dissection was performed to remove the peripheral and central nerve roots, resulting in only the sensory ganglia remaining. scRNA-seq experiments are the culmination of six independent bioreplicates. Sensory ganglia were dissociated in 40 units papain, 4 mg/ml collagenase, 10 mg/ml BSA, 1 mg/ml hyaluronidase, 0.6 mg/ml DNase in DMEM:F12 + 1% pen/strep + 12.5 mM glucose for 25 min at 37 °C. Digestion was quenched using 20 mg/ml ovomucoid (trypsin inhibitor), 20 mg/ml BSA in DMEM:F12 + 1% pen/strep + 12.5 mM glucose. Ganglia were gently triturated with fire-polished glass pipettes (opening diameter of approx. 150– 200 μm). Neurons were then passed through a 70-μm filter to remove cell doublets and debris. Neurons were pelleted and washed 4 times in 20 mg/ml ovomucoid (trypsin inhibitor), 20 mg/ml BSA in DMEM:F12 + 1% pen/strep + 12.5 mM glucose followed by 2× washes with DMEM:F12 + 1% pen/strep + 12.5 mM glucose all at 4 °C. After washing, cells were resuspended in 45 μl of DMEM:F12 + 1% pen/strep + 12.5 mM glucose.

### scRNA-seq analysis

Alignment, mapping, and general quality control was performed using the 10x genomics cell ranger pipeline. This pipeline generated the gene expression tables for individual cells used in this study. Briefly, ∼8000-10000 dissociated cells from DRG were loaded per 10x run (10x genomics chromium single cell kit, v3). Downstream reverse transcription, cDNA synthesis and library preparation were performed according to manufacturer’s instructions. All samples were sequenced on a NextSeq 550 with 58bp sequenced into the 3’ end of the mRNAs. As quality control filter, individual cells were removed from the dataset if they had fewer than 1,000 discovered genes, fewer than 1,000 unique molecule identifiers (UMIs) or more than 10% of reads mapping to mitochondrial genes. Non-neuronal cells were also removed from our analysis. Neuronal subtypes were identified as clusters using PCA/UMAP analysis combined with prior studies identifying marker genes for distinct DRG neuron subtypes. Non-neuronal cells were identified in one of two ways. In the first, if the cells contained prominent markers of endothelial or Schwann cells (Sox2, Ednrb) they were removed. In the second, if a cluster was clearly devoid of DRG sensory neuron markers (Pou4f1, Avil), they were also removed. Generally, we found approximately 20-25% of cells recovered showed signatures of non-neuronal identity based on these criteria. Differential gene expression analysis was performed on all expressed genes using the FindMarker function in Seurat using the Wilcoxon rank-sum test and a pseudocount of 0.001 was added to each gene to prevent infinite values. P values <10−322 were defined as 0, as the R environment does not handle numbers <10−322. Individual cell gene expression value were removed in the violin plots to allow for clarity due to the large number of cells

### AAV production and neonatal IP injection

AAV genome plasmids were constructed using standard cloning and molecular biology techniques. AAV (serotype 2/9) were packages and concentrated using transient transfection of pRC9, pHelpher and AAV-genome plasmid into 6-12 T225 flasks of HEK 293T cells. Viral containing supernatants were collected at 72 and 120 hours post-transfection. 120 hours post-transfection, cells were scraped off plates and also collected. AAV producing 293T cell pellets were extracted using a lysis buffer containing salt active nuclease (articzymes) in 40 mM Tris, 500 mM NaCl and 2 mM MgCl_2_ pH 8 (referred to as SAN buffer). Viral supernatants were concentrated via 8% PEG/500 mM NaCl precipitation and ultimately re-suspended in SAN buffer. Cleared viral lysates were then loaded onto a density gradient (optiprep) and subsequently concentrated using Amicon filters with a 100-kD molecular cutoff to a final volume of approximately 25-30uL. AAVs (2/9) were injected intraperitoneally (IP) into pups at P1-2. Pups were transiently anaesthetized by hypothermia and bevelled pipettes were used to deliver between 10^11^ and 10^12^ viral genomes in a volume of 10 μl (0.01% Fast Green, 1 × PBS). Skin, DRG, or spinal cord samples were collected from animals at least 3 weeks after AAV mediated transduction.

### Immunohistochemistry

We used a similar protocol as previously described ^28^. Animals were perfused with PBS then 4% PFA. Spinal column was dissected out and fixed in 4% PFA overnight (4°C). Hairy skin on the thigh or back was dissected out, treated with Nair Hair Removal Lotion and rubbed with tissue paper until all hair was removed. The skin was then washed and fixed in Picric acid-formaldehyde (PAF) fixative (Zamboni) overnight (4°C). After tissue fixation, the tissue was embedded in OCT (14-373-65, Fisher), frozen and stored in −80°C freezer. Upon cryosection, the tissue was sectioned with the thickness of 25 or 30 micron and mounted on the superfrost plus slides (12-550-15, Fisher). After drying in room temperature (RT) overnight, the slides were rehydrated and washed by PBS, and blocked by 5% Normal Donkey Serum for 2 hours at RT. Then primary antibody mix was added for 2 hours at RT or 1-2 days at 4°C. After washing the primary antibody with 0.02% PBS-Triton, the secondary antibody mixed was added for 2 hours at RT or 1-2 days at 4°C. At last, the slides were washed in PBS, mounted using DAPI Fluoromount-G (0100-20, Southern Biotech), and imaged using a confocal microscope (Zeiss LSM 700 or Zeiss LSM 900).

### *In situ* hybridization (RNAscope)

We used a similar protocol as previously described ^28^ following the recommended protocol from ACD bio website (https://acdbio.com/manual-assays-rnascope). DRGs were freshly dissected out of the animals, embedded in OCT and flash frozen. After cryosection at 25-micron thickness, the slides were fixed in 4% prechilled PFA in 4°C for 15 mins, then dehydrated using serial concentrations of ethanol. Next the sections were digested using Protease IV for 30 mins at RT. After washing, mix of RNAscope probes was added on the slide and incubated at 40°C for 2 hours, followed by adding and washing Amp1, Amp2, Amp3 and Amp4 sequentially according to the protocol. At last, the slides were mounted using DAPI Fluoromount-G (0100-20, Southern Biotech), and imaged using a confocal microscope (Zeiss LSM 700 or Zeiss LSM 900). Quantifications of the images were performed in ImageJ. Cells with clear fluorescent puncta were defined as positive for the corresponding RNA.

### Sparse labeling for single-neuron morphological analysis

For sparse labeling experiments, the available CreER lines were crossed to *Brn3a^cKOAP^* mice and the sparsity was controlled by tamoxifen dose. Aβ field-LTMRs or Aβ SA1-LTMRs were labeled using *TrkC^CreER^* treated with 0.01mg tamoxifen at P8. Aβ RA-LTMRs were labeled using *Ret^CreER^*treated with 0.03mg tamoxifen at E11.5. Aδ-LTMRs were labeled using *TrkB^CreER^*treated with 0.005mg tamoxifen at P5. C-LTMR were labeled using *TH^2A-CreER^*with 0.2mg tamoxifen at P21. *Sstr2^CreER-T^*^2a^*; Brn3a^cKOAP^* mice were given 0.1mg tamoxifen at P14. Sparse labeling of MRGPRD^+^ neurons relied on the leaky expression of *MrgprD^CreER^* without using tamoxifen.

For the sparse labeling in Cre lines, AAV9-FLEX-FLAP was injected into back hairy skin using reported procedure ^121^ or intrathecally as previously described ^122^. Back hairy skin injections were done in P9-P14 animals with a total volume of 2ul AAV injected into 4-6 spots across the back. Undiluted AAV (titer 1.2*10^13^) was used for *Cylstr2^Cre^*mice, while a 4X dilution was used for *Bmpr1b^Cre^* and *Smr2^Cre^*mice. Intrathecal (i.t.) injections were done in *MrgprB4^Cre^*or *MrgprA3^Cre^* mice that were 3-4 weeks old using 5 ul undiluted AAV9-FLEX-FLAP virus. For sparse labeling using *Trpm8^FlpO^* mice, 2ul of AAV9-fDIO-FLAP was injected into back hairy skin of *Trpm8^FlpO^*animals that were 9-12 days old. The animals were perfused 1-3 weeks after injection of tamoxifen or AAV.

### Whole-mount alkaline phosphatase staining of the skin and spinal cord

Whole-mount placental alkaline phosphatase staining protocol was adapted from a previous report ^121^. Animals of 4-6 weeks old were perfused in PBS then 4% PFA, then NAIR is applied on hairy skin on the back and thigh of the animals. After overnight fixation in Zamboni fixative (phosphate buffered picric acid-formaldehyde) at 4°C, tissue was washed in PBS and incubated at 65-68°C for 2-2.5 hours to inactivate the endogenous alkaline phosphatase activity, then washed with B3 buffer (0.1M Tris pH 9.5, 0.1M NaCl, 50mM MgCl_2_, 0.1% Tween-20) at room temperature. The substrate, NBT/ BCIP (3.4µL per ml of B3 buffer), was added to start the reaction. After 8-24 hours (depending on the expression level) at room temperature, tissue was washed in B3 buffer and pinned down in dishes before fixation in 4% PFA for one hour in room temperature. After fixation, dehydration started with putting the tissue 50% ethanol (1 hour), then 75% ethanol (1hour), at last 100% ethanol for 3 times, with overnight dehydration in the last time of 100% ethanol. After complete dehydration, tissue was cleared in BABB (Benzyl Alcohol: Benzyl Benzoate = 1:2) at room temperature for 30 mins. The tissue was then imaged under a Zeiss AxioZoom stereoscope. After imaging, the tissue was stored in 100% ethanol at 4°C.

### Quantification of skin arborizations in the sparse labeling experiments

The skin was flattened before imaging, and a Z-stack image was taken to cover the depth of the skin. Only arborizations that clearly belong to a single neuron were imaged. *Smr2^T^*^2a–Cre^ labeled neurons could also show lanceolate endings and circumferential endings although uncommon, which were excluded in the quantification because we focus on the SMR2/CGRP^+^ population that only showed free-nerve endings using *Smr2^T^*^2a–Cre^; *Calca^FlpE^* intersection (Figure S5). The arborization area was measured in ImageJ by drawing a polygon tightly around the skin arbors.

The axonal branch points were counted manually. Reconstruction of skin arborizations was done using SNT plugin ^123^.

### Axial level analysis of *Adra2a^T^*^2a–CreER^ labeling

Tamoxifen was administrated (i.p.) to *Adra2a^CreER^*; *Brn3a^cKOAP^* animals at P14 and P16 age, using a total dose of 4mg. Using *Brn3a^cKOAP^* reporter line avoids labeling of sympathetic fibers, which express Adra2a ^124^. Animals were perfused 2-3 weeks later. Cutaneous skin, various internal organs and spinal column were dissected out and fixed overnight at 4°C in either 4% PFA or Zamboni fixative. Spinal cord was dissected out carefully with most DRGs attached. The Whole-mount placental alkaline phosphatase staining protocol was the same as described above. After clearing with BABB, the DRGs attached to the spinal cord were imaged under a Zeiss AxioZoom stereoscope with Z-stack covering the depth of all DRGs. The L4/L5 DRGs were defined as the two large ganglion that connected to the two thickest nerves in lumbar level. The axial level of the other DRGs were based the relative distance (in the number of segments) to L4/L5. The number of labeled DRG cells were manually counted in ImageJ, although the dense labeling of certain DRGs (especially from T10-L1 level) and the resolution of the stereoscope would cause underestimation of cell numbers or render the image indiscernible. The cell counts from left and right DRGs of the same level were averaged for the same animals. There were occasional loss of DRGs during tissue processing, leading to variation of the number of datapoints for each axial level.

### *In vivo* epifluorescence calcium imaging

The mice of either sex at 3-6 weeks were anesthetized with inhalational isoflurane (1.8-2.3%) throughout the experiments. Body temperature was maintained at 37°C ± 0.5°C on a custom-made surgical platform. The back hair was shaved, and an incision was made over the lumbar vertebrae. Paravertebral muscles along L4 vertebral spine were dissected, then the bone on top of L4 DRG was removed by rongeur or by bone drill. Surgifoam sponges (Cat# 1972, Mckesson) and cotton was used to stop the bleeding. Custom-made spinal clamp was used to stabilize the spinal column.

The surgical preparation was then transferred to the platform under an upright epifluorescence microscope (Zeiss Axio Examiner) with 10X air objective (Zeiss Epiplan, NA=0.20). The light source is 470nm LED (M470L5, Thorlabs) with LED driver (LEDD1B, Thorlabs). A CMOS Camera (CS505MU1, Thorlabs) was triggered at 10 frames per second with a 50ms exposure time. All the recorded stimuli were synchronized with the camera and LED using a DAQ board (National Instrument, NI USB-6343).

### Stimuli applications

For assessment of mechanical thresholds using step indentation, receptive fields of a certain neuron within field of view was explored using a blunt probe. If a point gave the most robust calcium signal increase, the indenter was then placed on top of the point. Indentation was delivered using a mechanical stimulator (300C-I, Aurora Scientific) with a custom-made indenter tip with an end of ∼200um in diameter. The duration of indentation was 0.5 second for all experiments, except when examining the adaption properties, the indentation duration was increased to 3 seconds. The interval between indentation steps was set at least 6 seconds to allow the calcium signal to return baseline.

For assessment of polymodality, a square region (12.5 * 12.5 mm) on the mouse thigh was specified for all the following mechanical, electrical and thermal stimuli. Air puff (1PSI) was first delivered to the region. Then the stroke stimuli were delivered by cotton swab mounted vertically on a miniature load cell for force measurement (Model MBL, 50gram, Honeywell). The load cell was calibrated using standard weight. The force of stroke stimuli was maintained below 50mN. Next 6-gram, 10-gram and 26-gram Von Frey filaments (North Coast Medical) were applied on the same skin region sequentially. To fully cover the skin region for mechanical stimuli, a custom made Von Frey array was used. Individual 8-gram Von Frey filaments were made from Carbon Fiber (2004N11, McMaster Carr) by adjusting the length of the filament and indenting on a digital scale. Twenty-five of such filaments were mounted on a piece of acrylic (13*13 mm) in even distribution. The other side of the acrylic was connected to a manipulator (U-3C, Narishige). The last mechanical stimulation is pinch, which was delivered by a pair of cover glass forceps (11074-02, FST). A pair of FlexiForce sensor (A101, Tekscan) were mounted on the flat inner tip of the forceps, with rubber pads mounted on the sensor for better measurement. The force delivered by pinch were within 2-4 N range. Hair on the thigh was kept during air puff and stroke to preserve the natural condition, and shaved before using Von Frey filament and pinch stimuli for better spatial accuracy. All the mechanical stimuli were applied after 10 seconds of baseline recording, and lasted 20 seconds during which various spots across the selected skin region were stimulated intermittently.

Electrical stimuli were applied using custom-made bipolar electrode controlled by a Current Stimulator (DS3, Digitimer) triggered at 10 pulses per second. The stimulation intensity was no more than 1mA with a duration of 2ms for each pulse. The tip of bipolar electrode and skin was lighted dampened before the stimulation. The stimulation locations were changed within the selected skin region to cover the region, lasting no more than 30 seconds in total.

For thermal stimuli, a Peltier device (13*12*2.5mm, TE-65-0.6-0.8, TE technology) was controlled by a Temperature Controller (TEC1089/PT1000, Meerstetter Engineering) with an RTD Platinum (Pt) thermistor (2952-P1K0.161.6W.A.010-ND, Digikey) mounted on the surface of Peltier. A copper bar (12.7 *12.7 * 60mm) was mounted on the other side of the Peltier device using double-sided thermal tape, acting as a heatsink. The heatsink was connected to a manipulator (U-3C, Narishige) for positioning of the Peltier device. Before thermal stimuli, ∼50ul of thermal paste (Thermal Grizzly, Aeronaut) was evenly applied on the top of the Peltier surface, which was then gently pressed on the skin until good contact was formed, thus the thermal conductivity between the Peltier device and skin could be maximized. A thermocouple microprobe (IT-1E, Physitemp) was inserted between Peltier device and skin as a separate measurement of the applied temperature. Thirty seconds after contacting skin, temperature stimuli started from innocuous temperatures progressing to noxious temperatures, in the sequence of 35, 25, 20, 40, 15, 45, 10, 50, 5, 55°C. Each temperature step started with a baseline of 20 seconds holding at 32°C, then to the target temperature for 20 seconds, and back to the baseline (32°C) holding for 50 seconds before the next temperature step. The change rate for all temperatures was kept at 5°C s^-1^.

### Calcium Imaging analysis

For imaging analysis, motion correction and spatial high-pass filtering was conducted using a custom-written macro code that employed ImageJ plugin “moco” ^125^ and “Unsharp mask” filter, then regions of interest (ROIs) were manually picked. Cells with baseline signal and/or with calcium response were picked, and aligned across videos from various stimuli. The generated intensity measurement from ImageJ was further analyzed using MATLAB. In the calculation of ΔF/F, F was defined using baseline activity (average intensity before each stimulation). The mechanical threshold for each indentation session was determined by the first distinguishable calcium spikes aligned with step indentation. The amplitude of indentation step was determined by the maximum calcium peak (ΔF/F) within 0.5 seconds after the end of indentation step. The response amplitudes for air puff, stroke, Von-Frey filament and pinch were calculate as the maximum calcium response within the session. The warm/heat threshold was determined by the corresponding ascending temperature step where the first calcium spike occur, while the cooling/cold threshold was determined by the corresponding descending temperature step where the first calcium spike occur. The amplitude of thermal response was calculated as the maximum calcium peak aligned within each temperature step for the thermal sensitive cells.

For the polymodality plot (Figure 7), first we established three vectors for tuning to mechanical, heat and cold, respectively, each 120 degree apart with a distinct color vector. Then the maximal responses to mechanical, heat and cold from the same neurons were extracted. To determine the color of each point in the scatter plot, the maximal response to each modality was normalized to the summation of the maximal responses of all three modalities of each neuron. The dot product of the color vector and the corresponding normalized response vector gave the color of each dot. To determine the location of each point, log scale was applied to the magnitude of maximal response, to compress the range for visualization. The dot product of the direction vector of each modality and the corresponding log scale magnitude decided the location of each neuron in the scatter plot.

## Acknowledgements

We thank O. Mazor and P. Gorelik (HMS Research Instrumentation Core) for help with design and construction of the GCaMP imaging setup and stimulation devices, and Ginty lab and Sharma lab members for discussions and comments on the manuscript. We thank Caiying Guo and the Gene Targeting and Transgenics Facility at Janelia Research Campus for generating mouse lines. This work was supported by a Quan Predoctoral Fellowship (L.Q.), NIH grants NS097344 (D.D.G.), AT011447 (D.D.G.), 1DP2NS127278 (N.S.), The Klingenstein-Simons Foundation (N.S.), The Whitehall Foundation (N.S.), The Bertarelli Foundation (D.D.G.), The Hock E. Tan and Lisa Yang Center for Autism Research (D.D.G.), and the Lefler Center for Neurodegenerative Disorders (D.D.G.). D.D.G. is an HHMI investigator. This article is subject to HHMI’s Open Access to Publications policy. HHMI lab heads have previously granted a nonexclusive CC BY 4.0 license to the public and a sublicensable license to HHMI in their research articles. Pursuant to those licenses, the author-accepted manuscript of this article can be made freely available under a CC BY 4.0 license immediately upon publication.

## Author contributions

N.S., L.Q., and D.D.G. conceived the study. N.S. designed the new mouse lines, except for the *Cysltr2^T2A-Cre^* mouse line, which was designed and generated by T.V., M.P., V.K.K., and I.C. L.Q. N.S. and M.I. characterized the mouse lines and performed all histological and morphological analyses with help from D.S, C.W, K.L and P.R. L.Q. performed the GCaMP imaging experiments, with help from N.S. N.S., L.Q., and D.D.G. wrote the paper with input from all authors.

## RESEOURCE AVAILABILITY

### Lead contact

<u>Further</u> information and requests for resources and reagents should be directed to and will be fulfilled by the lead contact, David Ginty.

### Materials availability

Requests for mouse lines generated in this study should be directed to and will be fulfilled by David Ginty.

### Data and code availability

All data reported in this study will be shared by the lead contact upon request. All original code is available in this paper’s supplemental information.

Any additional information required to reanalyze the data reported in this paper is available from the lead contact upon request.

### Key resources table

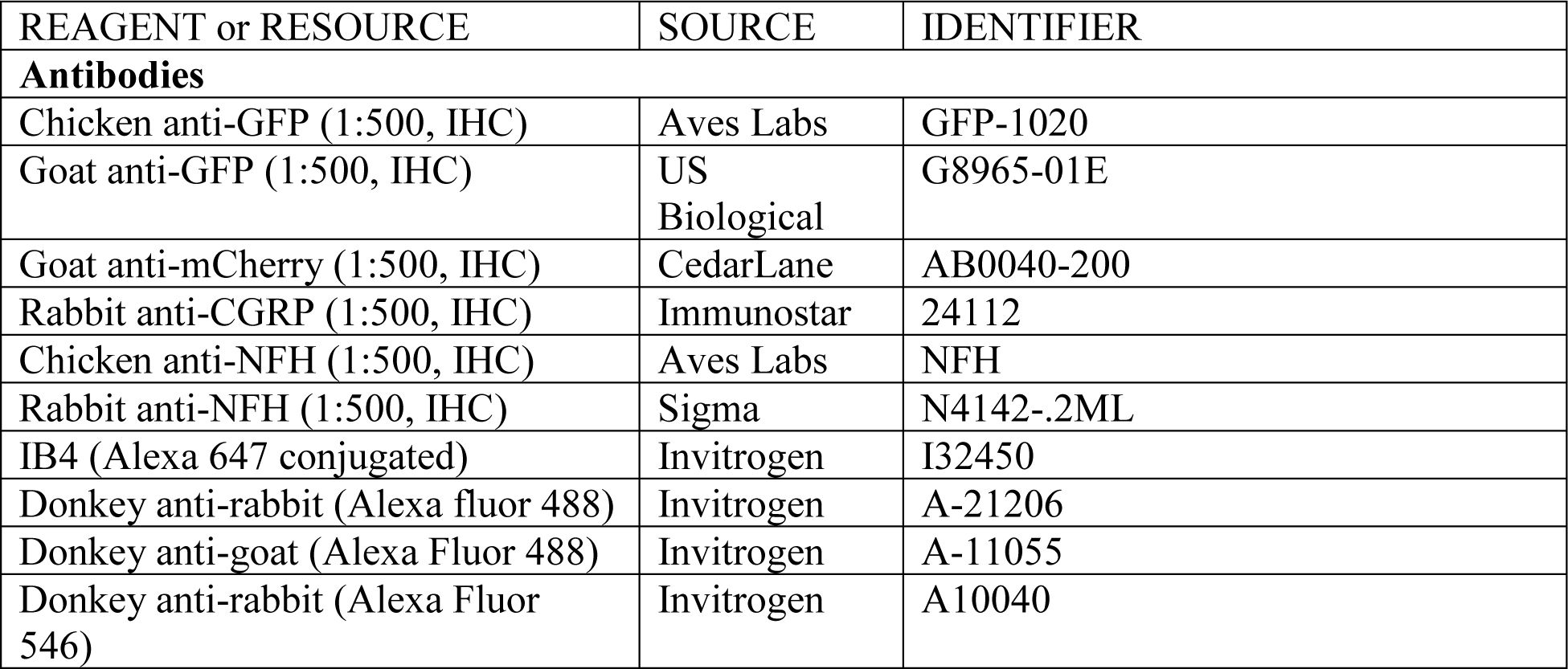
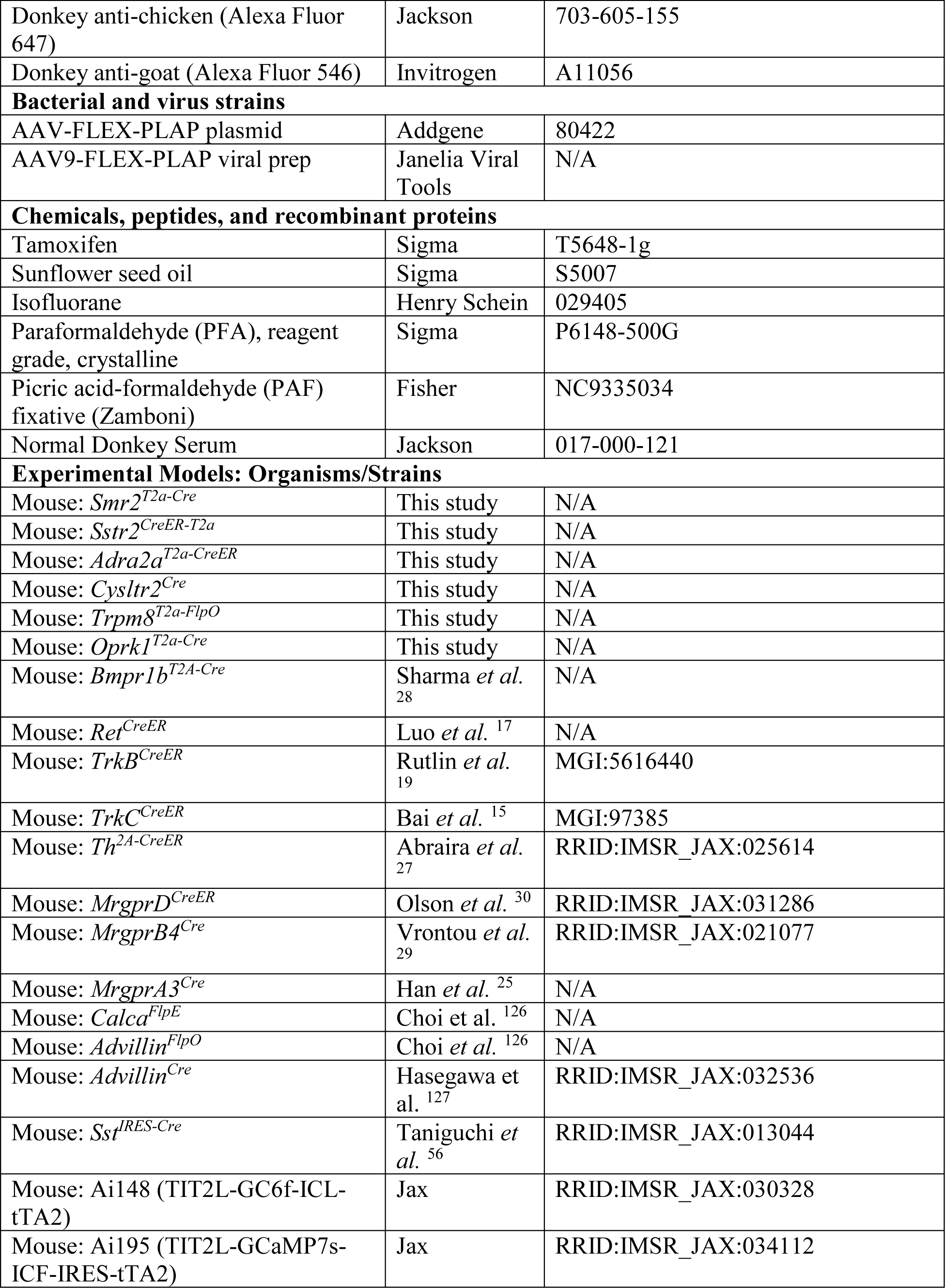
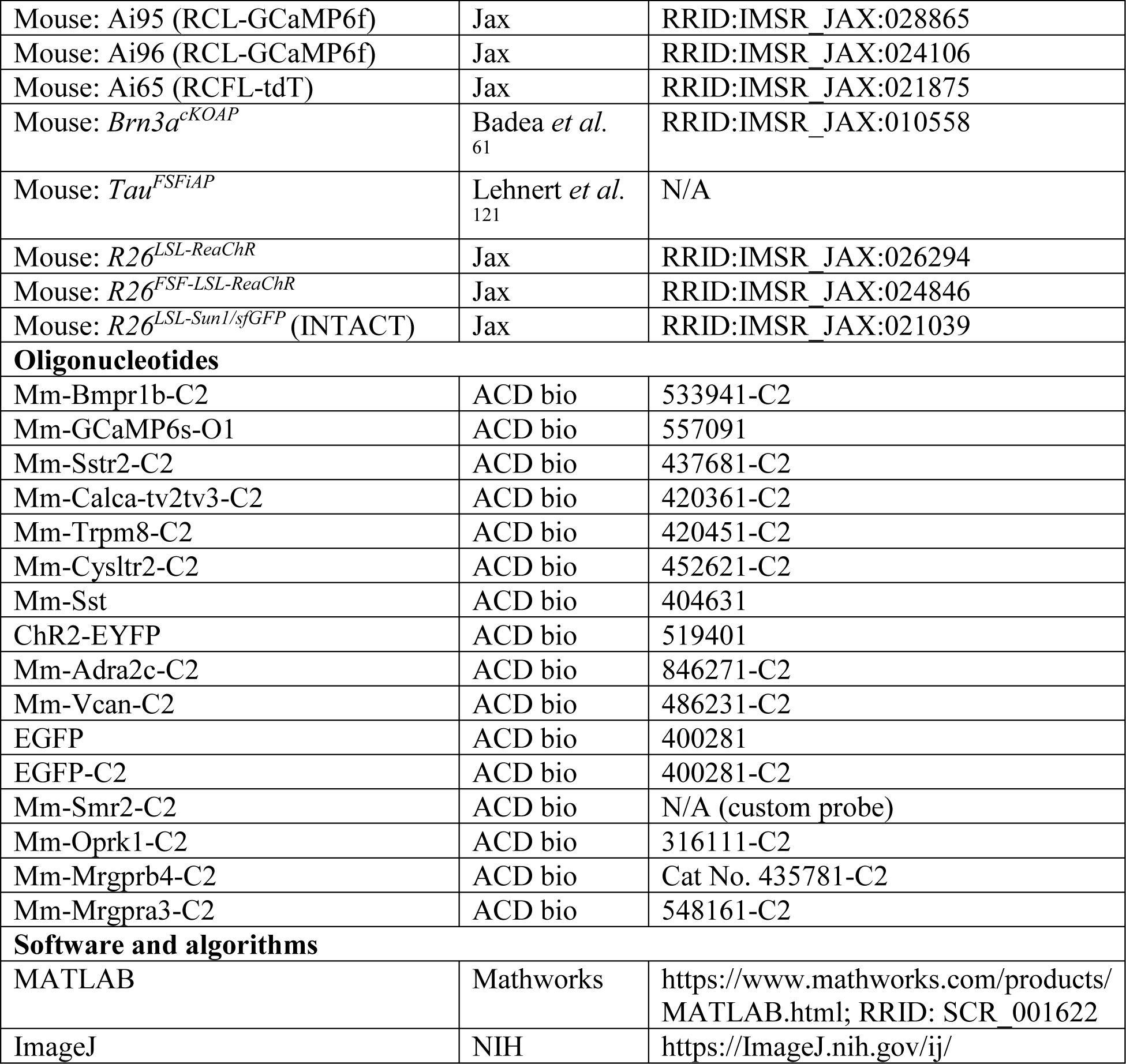

## Supplementary Figure Legends

**Figure S1.**
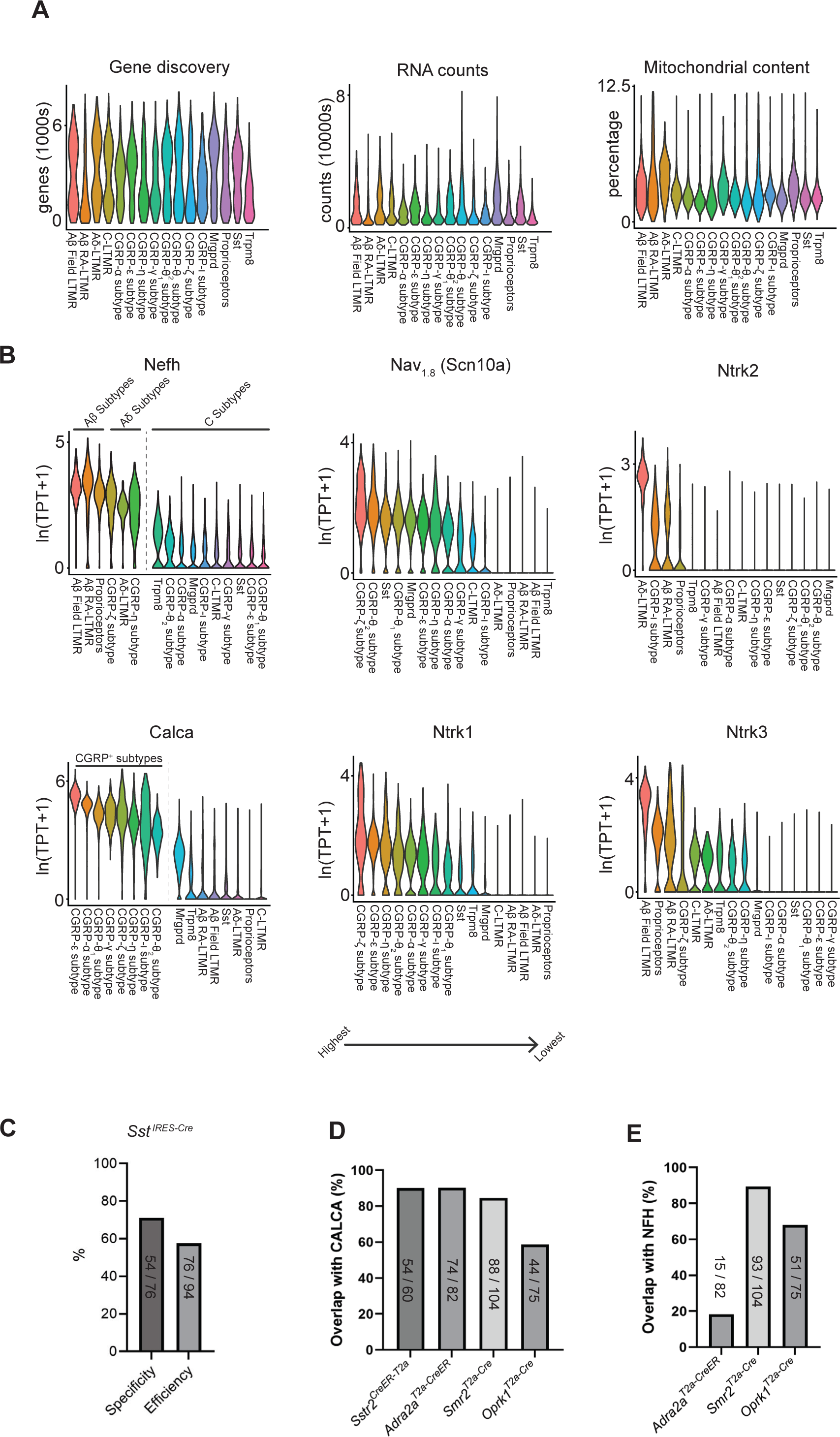
Additional information from the scRNA-seq dataset and further characterization of the new genetic tools. **(A)** Additional QC metrics and descriptive statistics for the scRNA-seq data. **(B)** The expression of common marker genes in transcriptionally defined DRG subtypes. **(C)** The specificity and efficiency of labeling by *SST^IRES-Cre^* mice crossed to *R26^LSL-Sun^*^1^*^/sfGFP^* mice determined by RNAscope. Specificity and efficiency was defined as described in Figure 1D. **(D-E)** The overlap with Calca (D) and NFH (E) for DRG neurons labeled using new genetic tools and examined by RNAscope.

**Figure S2.**
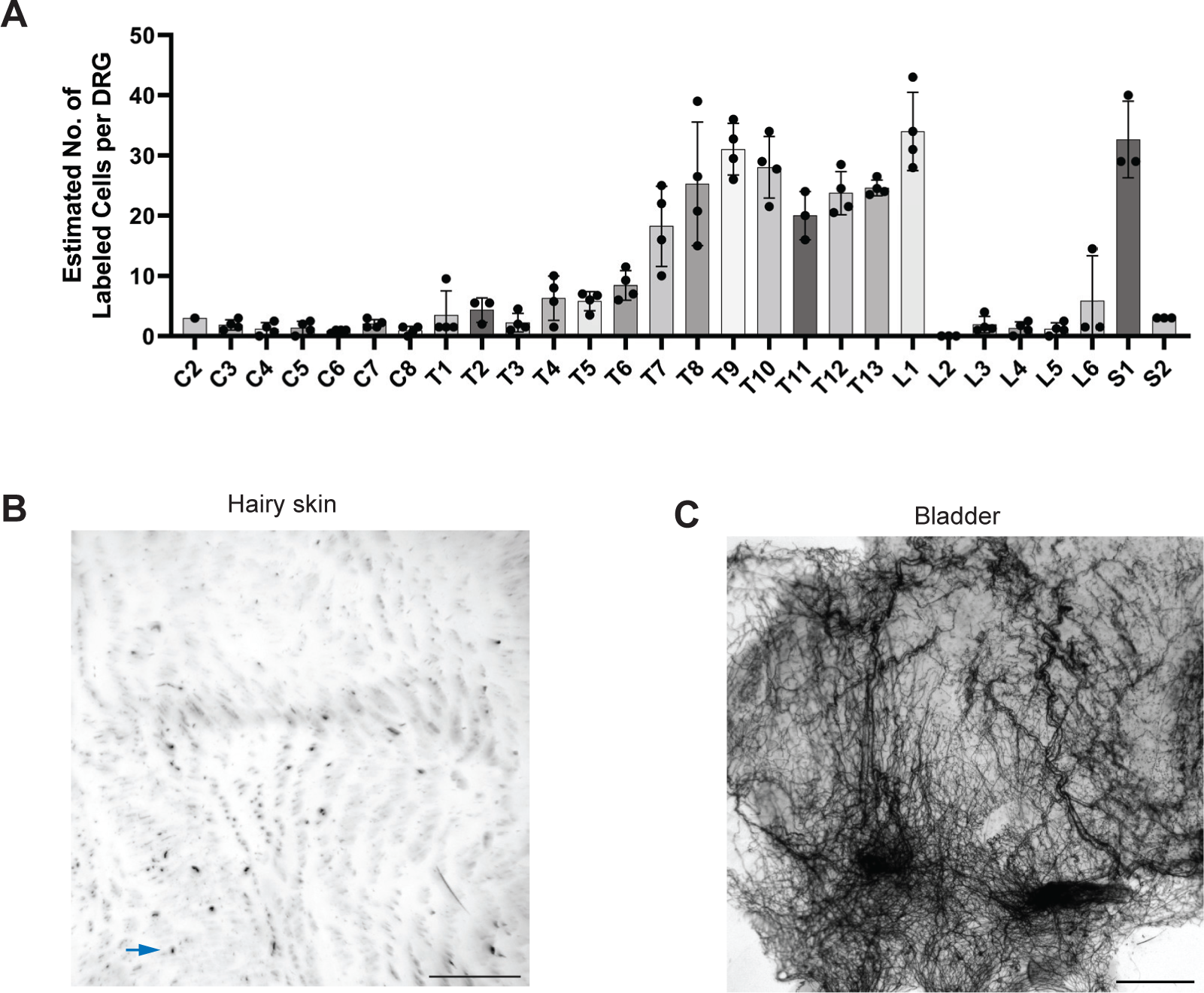
*Adra2a*^T2a–CreER^ labeled neurons (CGRP-γ) innervate internal organs but not the skin. **(A)** Estimated number of labeled DRG neurons across different axial levels using *Adra2a*^T2a–CreER^; *Brn3a^cKOAP^* mice and whole mount AP staining of spinal cords with DRGs attached. Each data point is the averaged number between left and right DRG of the same axial level from the same animal (4 animals in total). DRGs that were lost or that were too dense to count in the whole mount staining were not included. **(B)** Examples of whole mount AP staining of back hairy skin. The arrow points to the typical background signal. Only a total of 1-5 cutaneous neurons were identified from each animal. **(C)** Example of whole mount AP staining of the bladder **(C)**. Scale bar in **B** and **C** is 1mm.

**Figure S3.**
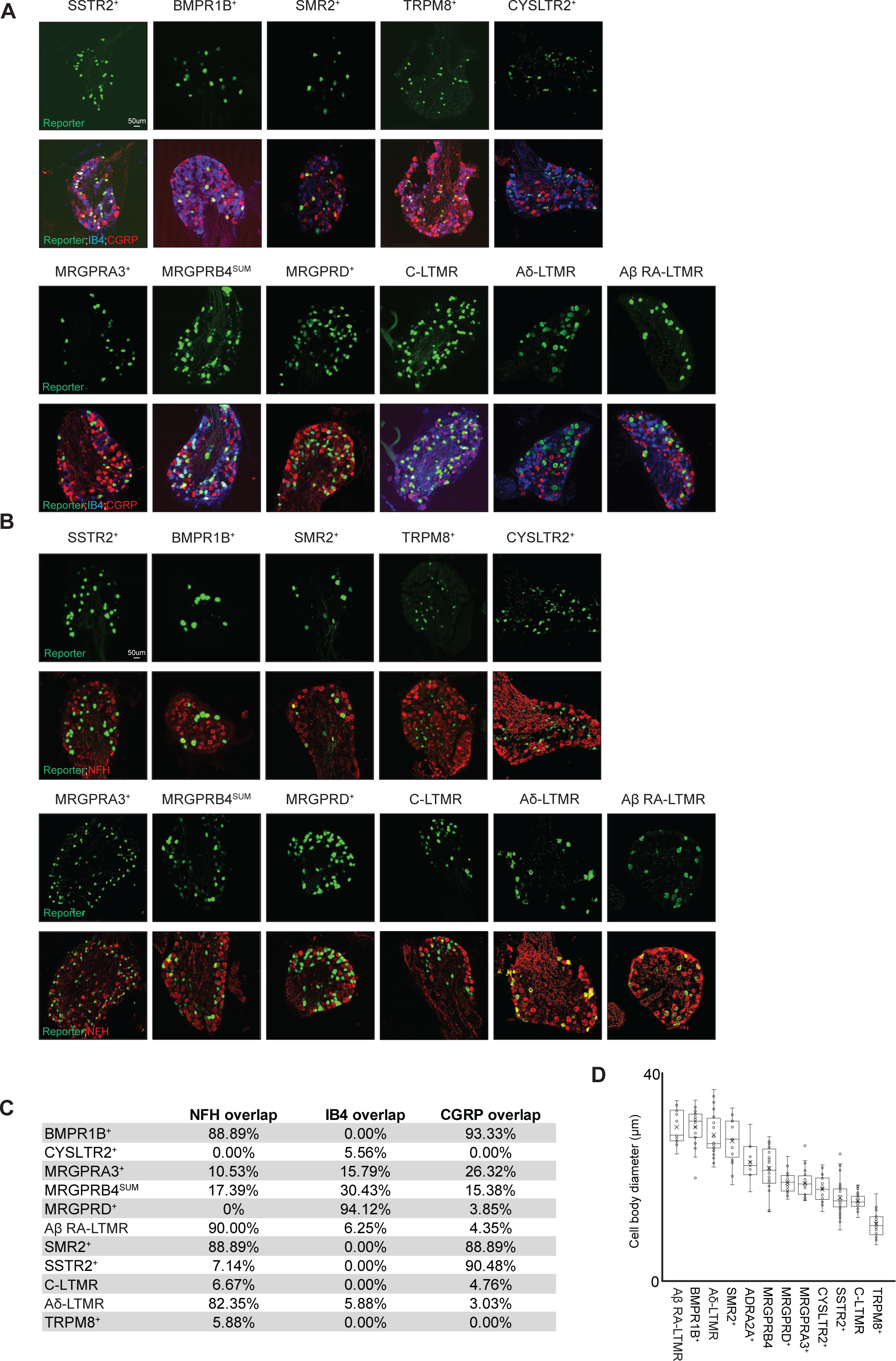
Immunohistochemical analysis of DRG sensory neuron subtypes. **(A)** DRG sections obtained from the mice harboring the new recombinase alleles (top row) and previously established alleles (bottom row) were co-stained with the reporter, CGRP, and IB4. **(B)** DRG sections obtained from the new recombinase alleles (top row) and previously established alleles (bottom row) were co-stained with the reporter and NFH. **(C)** Quantification of the percent overlap of labeled cells with respect to CGRP **(A)**, IB4 **(A)** and NFH **(B)**. **(D)** Quantification of the cell body diameter across the DRG subtypes. All scale bars are 50 µm.

**Figure S4.**
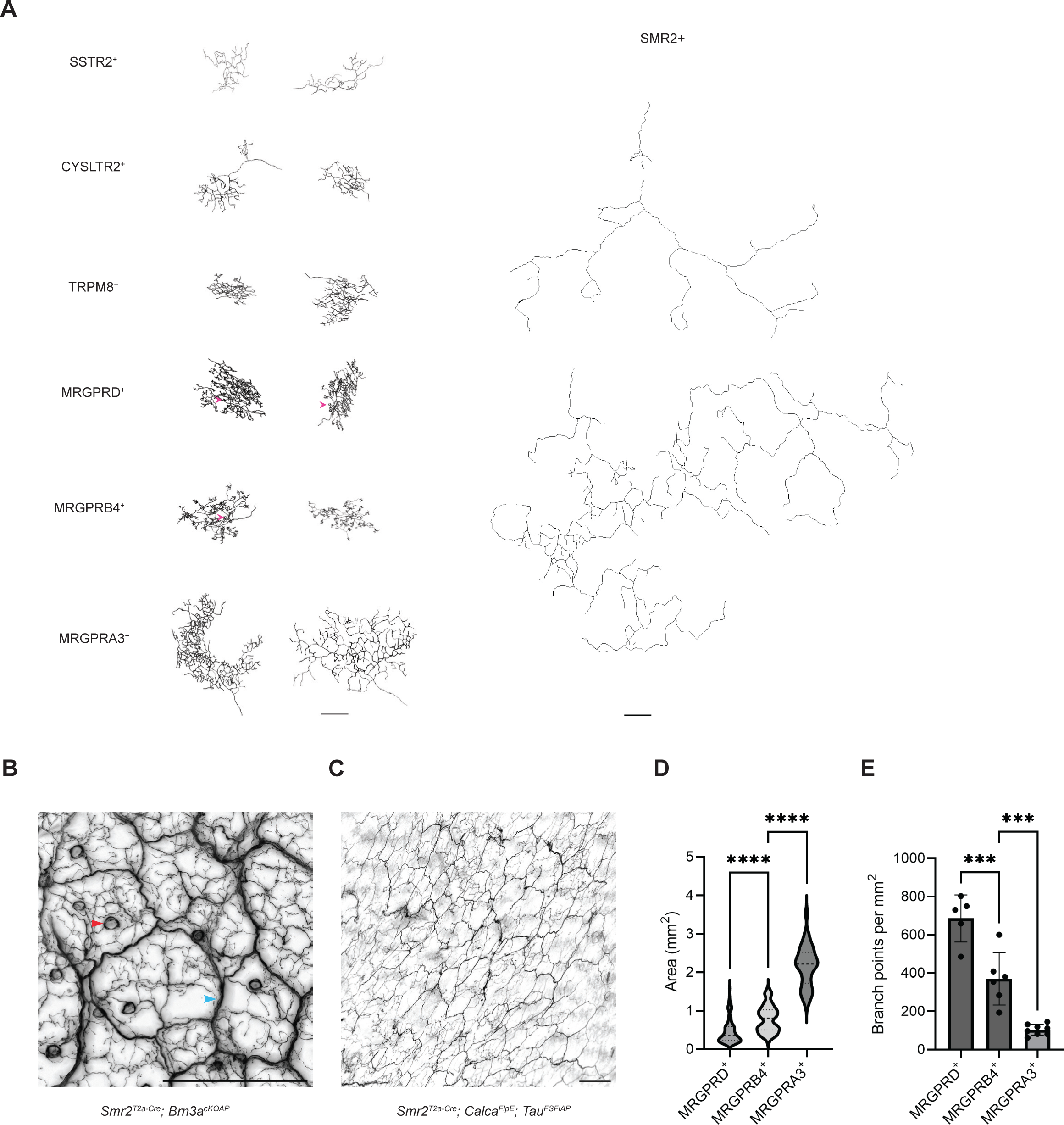
Morphological diversity of “free-nerve endings”. **(A)** Additional reconstructed examples of individual “free-nerve ending” neurons across the different populations. Arrows in magenta point to the “circumferential-like” terminals of MRGPRD^+^ and MRGPRB4^+^ neurons. **(B)** *Smr2^T^*^2a–Cre^; *Brn3a^cKOAP^* occasionally labels circumferential endings (red arrow). Blue arrow points to the thick nerve fiber under the epidermis. **(C)** Intersection genetic labeling strategy using *Smr2^T^*^2a–Cre^; *Calca^FlpE^*; *Tau^FSFiAP^* mice is specific for labeling CGRP-ζ (SMR2^+^) populations, only showing free-nerve endings. **(D-E)** Comparison of the arborization area **(D)** and branching density **(E)** for MRGPRD^+^, MRGPRB4^+^, and MRGPRA3^+^ neurons. One-way ANOVA test, ****p<0.0001, ***p<0.001. All scale bars are 500 µm.

**Figure S5.**
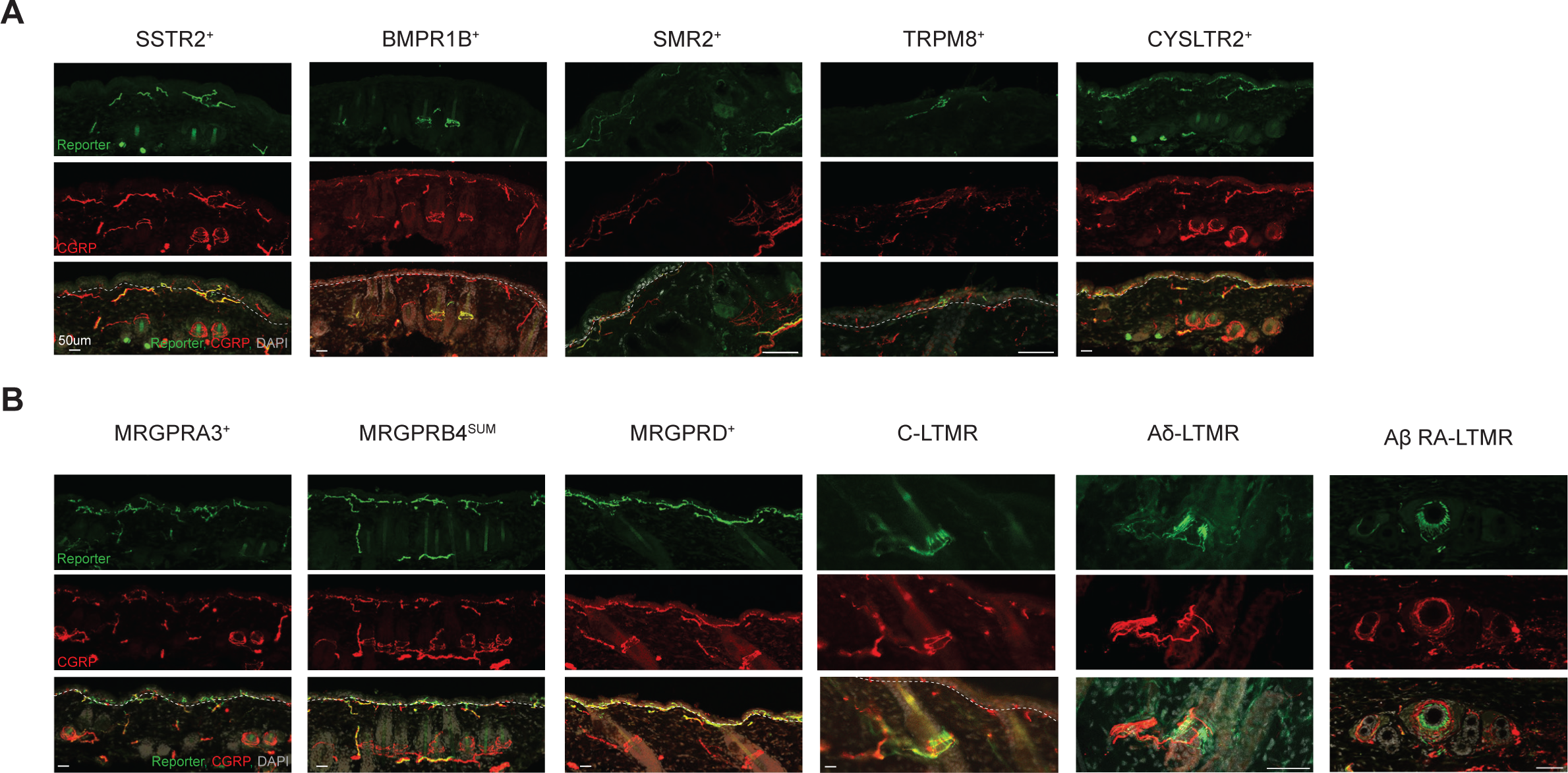
Hairy skin innervation patterns of DRG sensory neuron subtypes. **(A-B)** Representative immunostaining images of hairy skin samples from new driver lines and existing genetic tools. The panels show the staining of the reporter (GFP or tdTomato) (top), CGRP (middle), and the merge of the two overlaid with DAPI (bottom). The white dashed line indicates the border between the epidermis and dermis.

**Figure S6.**
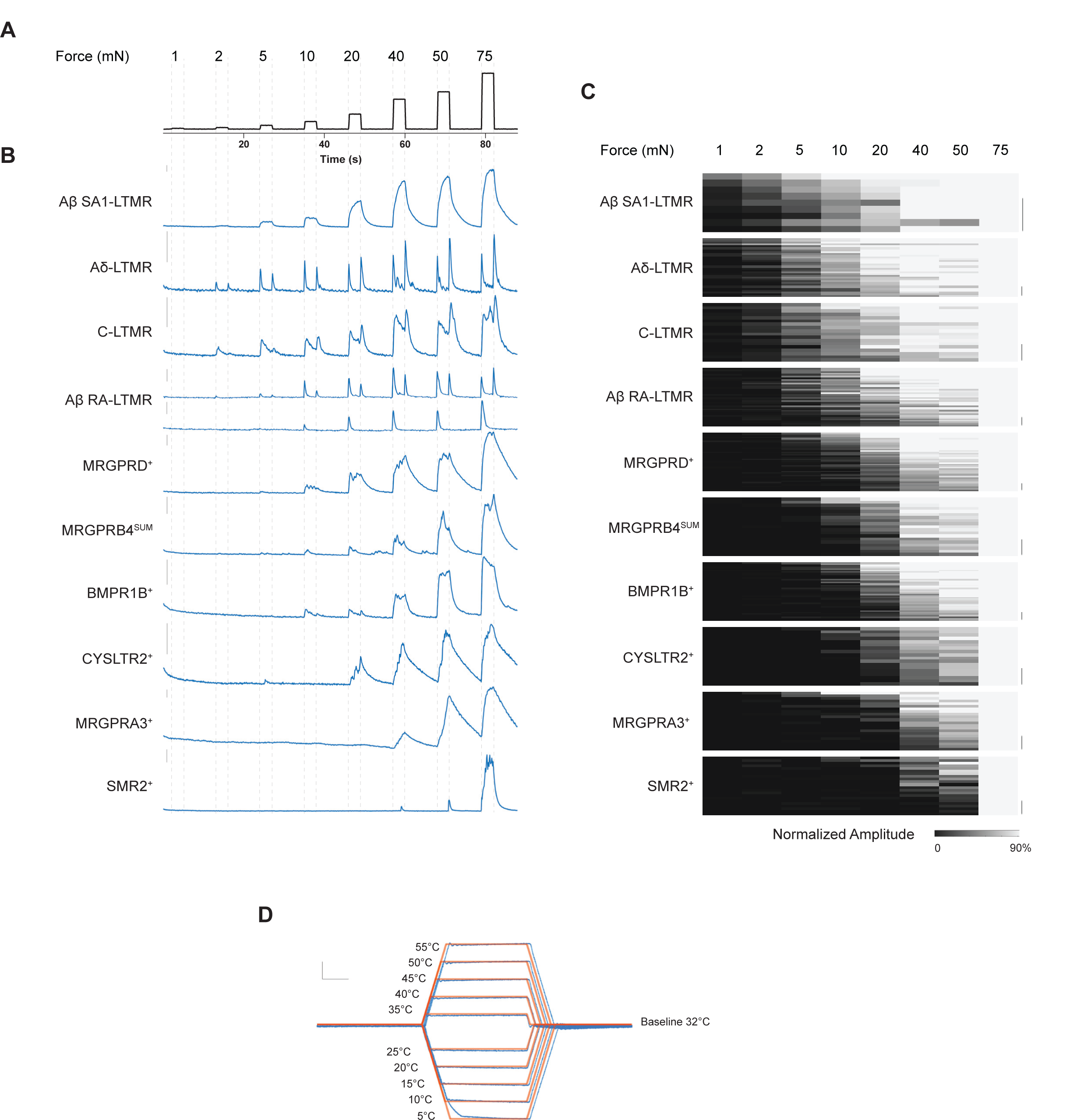
Adaption properties and stimulus-response relationships of DRG neurons subtypes. **(A)** Prolonged indentation steps, ranging from 1mN to 75mN. The duration of each step indentation for these experiments is 3 seconds. **(B)** Representative response of each DRG neuron subtype aligned to the indentation stimuli. Scale bar is 20% ΔF/F. **(C)** Normalized amplitudes of individual neurons from each DRG neuron subtype in response to skin indentation (duration 0.5 second). Each row represents a neuron. The amplitude of calcium response is normalized to the response at 75mN. The vertical scale refers to 5 cells. **(D)** Temperature stimuli used for the polymodality assay. Red lines refer to the command temperature, and blue lines represent the recorded temperature between the Peltier device and skin. The temperature steps are applied in the following order: 35, 25, 20, 40, 15, 45, 10, 50, 5, and 55°C. The rate of temperature change was kept at 5°C s^-1^. Scale bars are 5 °C (Y axis) and 5 seconds (X axis).

**Figure S7.**
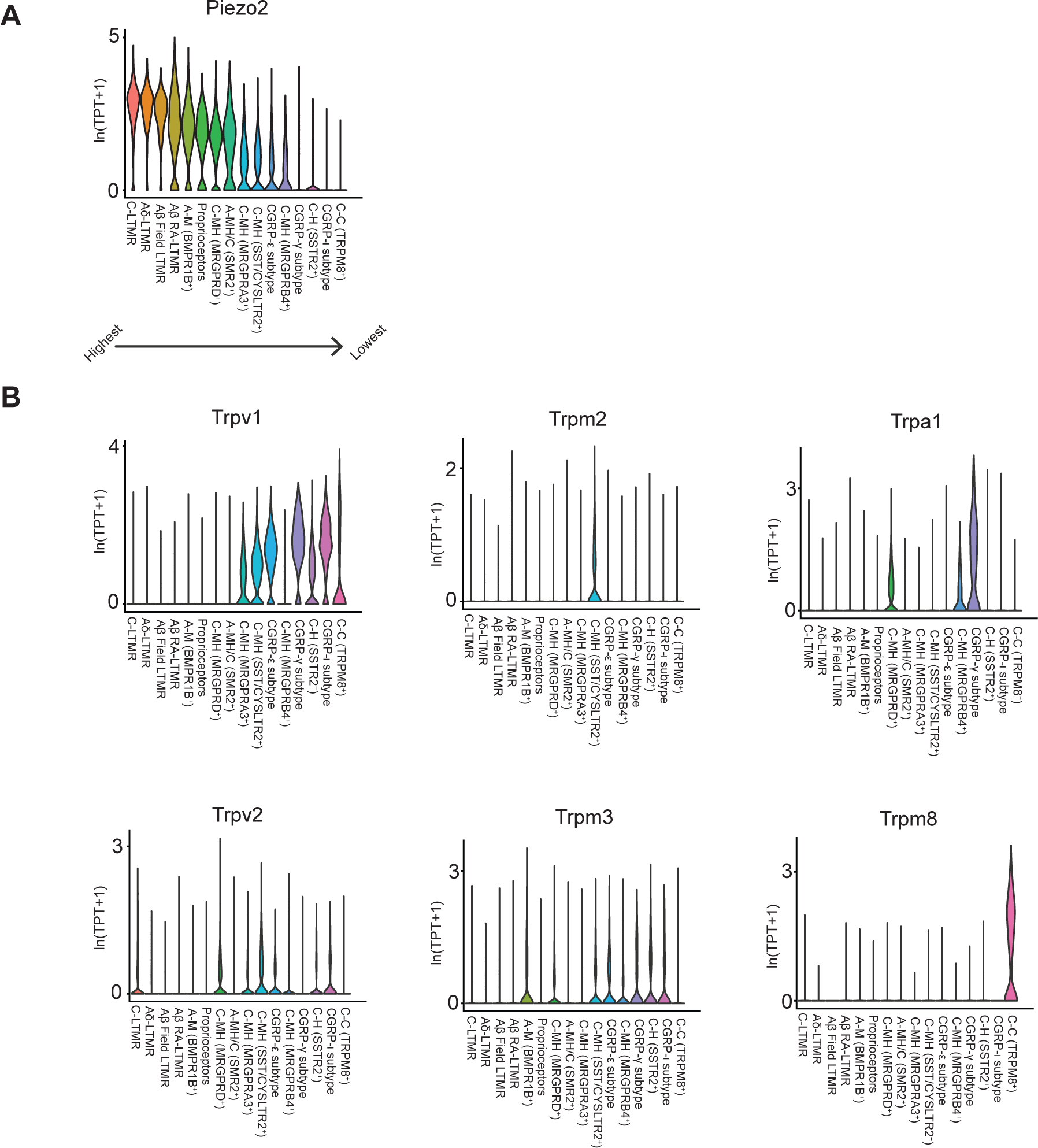
Expression profiles of Piezo channels and thermo-TRP channels across DRG neuron subtypes. **(A)** Violin plots displaying expression profile of Piezo channels. The DRG subtypes are sorted based on their relative level of expression of Piezo2 (highest left, lowest right). **(B)** Expression patterns of thermoTRP channels for genes with significant levels of expression in the DRG. TPT: tags per ten thousand.

